# Assessment of the relationships between agroecosystem condition and soil erosion regulating ecosystem service in Northern Germany

**DOI:** 10.1101/2020.05.26.116285

**Authors:** Paula Rendon, Bastian Steinhoff-Knopp, Philipp Saggau, Benjamin Burkhard

**Affiliations:** Institute of Physical Geography & Landscape Ecology, Leibniz University of Hannover, Hannover, Germany; Institute of Geography, Christian Albrechts University, Kiel, Germany; Leibniz Centre for Agricultural Landscape Research (ZALF) Müncheberg, Germany

**Author notes:** Corresponding author (PR).

**Keywords:** soil, ecosystem state, ecosystem condition indicators, control of erosion rates, Lower Saxony, agriculture, pressures.

## Abstract

Ecosystems provide multiple services that are necessary to maintain human life and activities. Agroecosystems are very productive suppliers of biomass-related provisioning ecosystem services, e.g. food, fibre and energy. At the same time, they are highly dependent on respective ecosystem condition and regulating ecosystem services such as soil fertility, water supply or soil erosion regulation. Assessments of this interplay of ecosystem conditions and services are very important to understand the relationships in highly managed systems. Therefore, the aim of this study is twofold: First, to test the concept and indicators proposed by the European Union Working Group on Mapping and Assessment of Ecosystems and their Services (MAES) for the assessment of agroecosystem condition at a regional level. Second, to identify the relationships between ecosystem condition and the delivery of ecosystem services. For this purpose, we applied an operational framework for integrated mapping and assessment of ecosystems and their services. We used the proposed indicators to assess the condition of agroecosystems in Northern Germany and the provision of the regulating ecosystem service control of erosion rates. We used existing data that are available from official databases for the calculation of the different indicators. We show maps of environmental pressures, ecosystem condition and ecosystem service indicators for the Federal State of Lower Saxony. Furthermore, we identified areas within the state where pressures are high, conditions are limited, and more sustainable management practices are needed.

Despite the limitations of the indicators and data availability, our results show positive, negative and no significant correlations between the different pressures and condition indicators, and the control of erosion rates. Although the idea behind the MAES framework is to show the general condition of an ecosystem, when looking at the relationships between condition and ecosystem services, we identified that not all the indicators - as they are proposed- are suitable to explain to what extent ecosystems are able to provide certain ecosystem services. Further research on other ecosystem services provided by agroecosystems would facilitate the identification of synergies and trade-offs. Moreover, the definition of a reference condition, although complicated for anthropogenically highly modified agroecosystems, would provide a benchmark to compare information on the condition of the ecosystems, leading to better land use policy and management decisions

## 1. Introduction

Human well-being and activities are strongly dependent on ecosystems, their biodiversity, condition, functionality and capacity to deliver multiple services. Ecosystem condition entails a clear link to ecosystem services and has been defined as the overall quality of an ecosystem unit in terms of its main characteristics that underpin its capacity to generate ecosystem services [1]. Assessing ecosystem condition can help to understand to what extent ecosystems are able to provide services in a sufficient quantity and quality. Some studies have focused on the links between natural capital and ecosystem services [2] and the relationships between biodiversity and ecosystem services [3]. However, understanding the relationships between ecosystem condition and the provision of ecosystem services is a remaining research gap [4, 5].

Ecosystem condition has been integrated in international and European Union (EU) policies for setting up sustainability and conservation targets and comprises concepts such as ecosystem state, quality, status, health, integrity and functioning [6]. The Sustainable Development Goals (SDGs), for instance, were adopted by the Member States of the United Nations in 2015, as “a call for action to end poverty, protect the planet and improve the lives and prospects of everyone everywhere” [7]. In particular, for biodiversity, goal 15 aims to protect, restore and promote sustainable use of terrestrial ecosystems, sustainably manage forests, combat desertification, and halt and reverse land degradation and stop biodiversity loss.

In 2011, the EU adopted the Biodiversity Strategy to 2020 [8] and established six targets to halt the loss of biodiversity and ecosystem services in the EU by 2020. These targets are oriented towards the protection of species and habitats, the maintenance and restoration of ecosystems, sustainable agriculture and use of forests, sustainable fishing and healthy seas, fighting invasive alien species and stopping the loss of biodiversity. The 7^th^ Environmental Action Programme (EAP) was adopted by the EU in 2013 [9] and it reinforces the targets and actions of the Biodiversity Strategy. The EAP aims to protect natural capital, stimulate resource-efficient, low-carbon growth and innovation, and safeguard human health and well-being, while respecting Earth’s limits [9]. Some topics that have been highlighted in both policies and that need further action at EU and national level are the protection of soils and the sustainable use of land, as well as forest resources.

The EU has established a dedicated working group on Mapping and Assessment of Ecosystems and their Services (MAES) [10] to support the implementation of Action 5 of Target 2 of the EU Biodiversity Strategy to 2020. This action requires that all Member States of the EU map and assess the state of ecosystems and their services in their territory, assess the economic value of such services and integrate these values into accounting and reporting systems at the national and EU level [8]. As part of its work, MAES has developed a conceptual framework [11] and suggested a series of indicators to assess the condition of different ecosystem types including agroecosystems in the EU [12]. However, these indicators still need to be tested at EU, national/sub-national and regional level; and links with ecosystem services still need to be further investigated. Our approach in this study is to test the indicators suggested by MAES for environmental pressures, ecosystem condition and the relationships with ecosystem services on a regional level, specifically in agroecosystems in Northern Germany, focusing on the soil erosion narrative.

Agroecosystems account today for almost half of the area of land use in the EU [13]. In Germany specifically, more than half of the surface area is used for agriculture [14]. Agricultural land provides, on the one hand, multiple ecosystem services, especially biomass-related services such as food, fibre, fodder or energy, which are essential for the maintenance of quality of life [15, 16]. On the other hand, agriculture itself is strongly dependent on ecosystem services such as nutrient regulation, water supply, pollination (for selected crop species) or soil erosion regulation [17, 18]. Changes in the condition of agroecosystems impair the availability of these services and hence jeopardise human well-being. Environmental pressures such as soil erosion, soil biodiversity decline, soil compaction, organic matter decline, soil sealing, and contamination, together with changing climate and water regimes, degrade these ecosystems [19]. In this regard, the maintenance of good condition of agroecosystems is essential to guarantee their resilience, halt biodiversity loss and preserve the sustainable provision of multiple ecosystem services.

Agroecosystems are strongly modified semi-natural systems that are managed with the purpose of enhanced ecosystem service delivery with a strong focus on provisioning services [20]. These ecosystem service outputs are, at least in conventional farming, strongly based on substantial anthropogenic human system inputs including fertilizer, insecticides, herbicides, energy, labour and machinery use and in some cases also irrigation water [21]. Besides ecosystem service outputs, agroecosystem service delivery has also significant environmental effects in the form of externalities such as greenhouse gas emissions, biodiversity loss or water eutrophication [18].

Due to the often long-term human interferences in these systems, there are shortcomings when defining a (natural) reference condition of agroecosystems. Agroecosystem condition cannot only be based on the physical and ecological properties of plants and soils, but must take into consideration human interferences that are characteristic of agroecosystems [16]. Agroecosystems are considered to be in good condition when they support biodiversity, when abiotic resources such as water, soil and air are not depleted and when they supply multiple provisioning, regulating and cultural ecosystem services [12]. Nevertheless, the establishment of threshold values to determine whether an agroecosystem is in good or bad condition is still under debate [22]. There is no explicit time reference condition as, for example, before the industrial revolution or other reference times like in other ecosystem types. Besides these temporal issues, also the involvement of multiple stakeholders (farmers, policy makers, planners, consumers, environmental groups), which may have different interests and perceptions about the condition of agroecosystems [23], hampers the reference state definition.

This study focuses on the ecosystem service *control of erosion rates* and takes into account that soil erosion by water is a major problem in soil conservation in the EU [24, 25]. Soil erosion by water accounts for the largest share of soil loss in Central European agricultural ecosystems, especially in areas with steep slopes [26]. Unsuitable management activities threaten croplands by increasing the vulnerability of soils to erode [24]. The objective of this study is to conduct an integrated assessment of ecosystems for one exemplary ecosystem service in a specific ecosystem type and by making use of the indicators proposed by MAES for the assessment of ecosystem condition. The study applies an operational framework for integrated mapping and assessment of ecosystem and their services suggested by Burkhard et al [27] in agroecosystems in the Northern German Federal State of Lower Saxony. For this purpose, a stepwise approach is followed, including: i) the identification of the policy objective „healthy soils“ (in our study exemplified by soil erosion regulation) as a theme for the assessment; ii) the identification and mapping of agroecosystems; and iii) the selection, quantification and mapping of indicators of agroecosystem condition.

The main aim of this study is to test the feasibility of the indicators proposed by MAES for the assessment of agroecosystem condition at a regional level. Thereby, we hope to contribute to the further methodological development and to increase the applicability of the MAES framework and indicators, rather than focussing on obtaining the most thorough and accurate results for a case study. The experience gained with the indicator application will be relevant for other (also non EU/MAES-related) comparable indicator-based studies in agroecosystems, their condition and ecosystem services.

The article is organized as follows: First, we explain the methodological approach used for the integrated assessment. Then we present the evaluation of the environmental pressures, ecosystem condition and the ecosystem service *control of erosion rates* by showing maps of the different indicators. We also statistically analyse the relationships between ecosystem condition and the *control of erosion rates*. Then we discuss the main limitations of the indicators and the connection between environmental pressures, ecosystem condition and ecosystem services and conclude with recommendations for further improvement of the MAES framework and indicators.

## 2. Methods

### 2.1. Study area

Lower Saxony is a federal state located in the northwest of the Federal Republic of Germany adjoining the North Sea. The federal state has an area of 47,620 km^2^ (Fig 1). Its climate is characterized as sub-oceanic with average temperatures ranging from 8.3 to 9.5°C and mean precipitation averages ranging from 654 mm in the south-west to 840 mm in the central and northern parts [28]. Agriculture is the main land use in Lower Saxony with 2.6 million hectares (approximately 53.7% of the total territory), of which 1.9 million hectares are arable land, 0.7 million hectares are permanent grassland and around 20,000 hectares are permanent crops [29].

**Fig 1.**
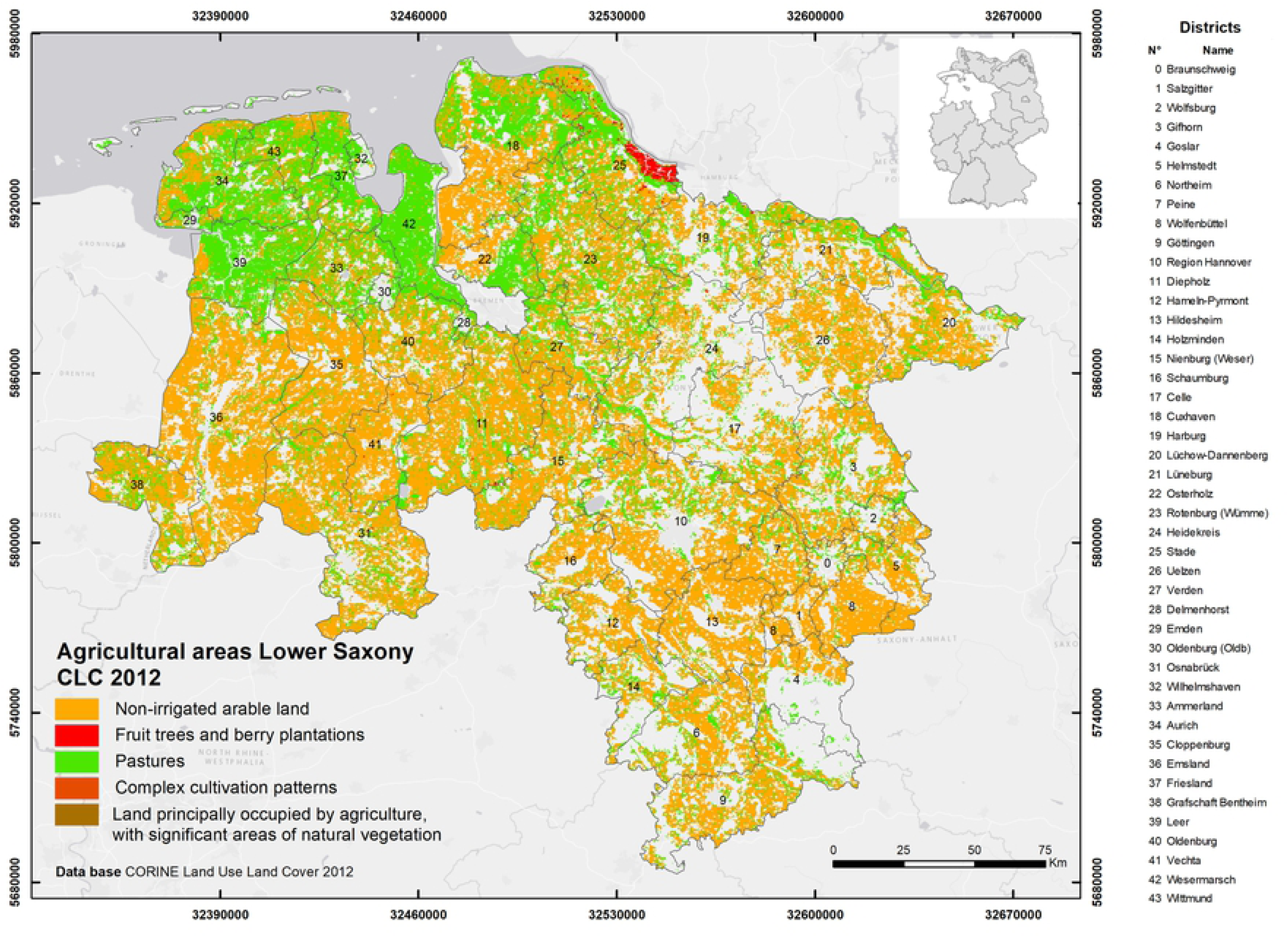
Agricultural areas in Lower Saxony (data source: CORINE Land Use Land Cover, 2012)

More than half of the arable land in Lower Saxony is used to grow cereals (mainly winter wheat and winter barley), the remaining part for fodder crops, oilseeds, or potatoes. The farm sizes are very diverse and range from a few hectares of specialized horticultural businesses to large arable farms with several hundred hectares. On average the farms have a size of 83 ha and about three quarters of all farms keep animals, especially dairy cattle and pigs. Farm type, specialization and size can be interpreted as a function of soil fertility, climate conditions and historical land use strategies [29].

### 2.2. Conceptual framework

In this study, we used the operational framework that guides integrated mapping and assessment of ecosystems and their services proposed by Burkhard et al [27]. Fig 2 provides a graphical summary of the framework with its nine steps: (1) theme identification; (2) identification of ecosystem type; (3) mapping of ecosystem type; (4) definition of ecosystem condition and identification of ecosystem services to be delivered by agroecosystems; (5) selection of indicators for ecosystem condition and ecosystem services; (6) quantification of ecosystem condition and ecosystem services indicators; (7) mapping ecosystem condition and ecosystem services; (8) integration of results; and (9) dissemination and communication of results. These steps are described in detail in the next paragraphs.

**Fig 2.**
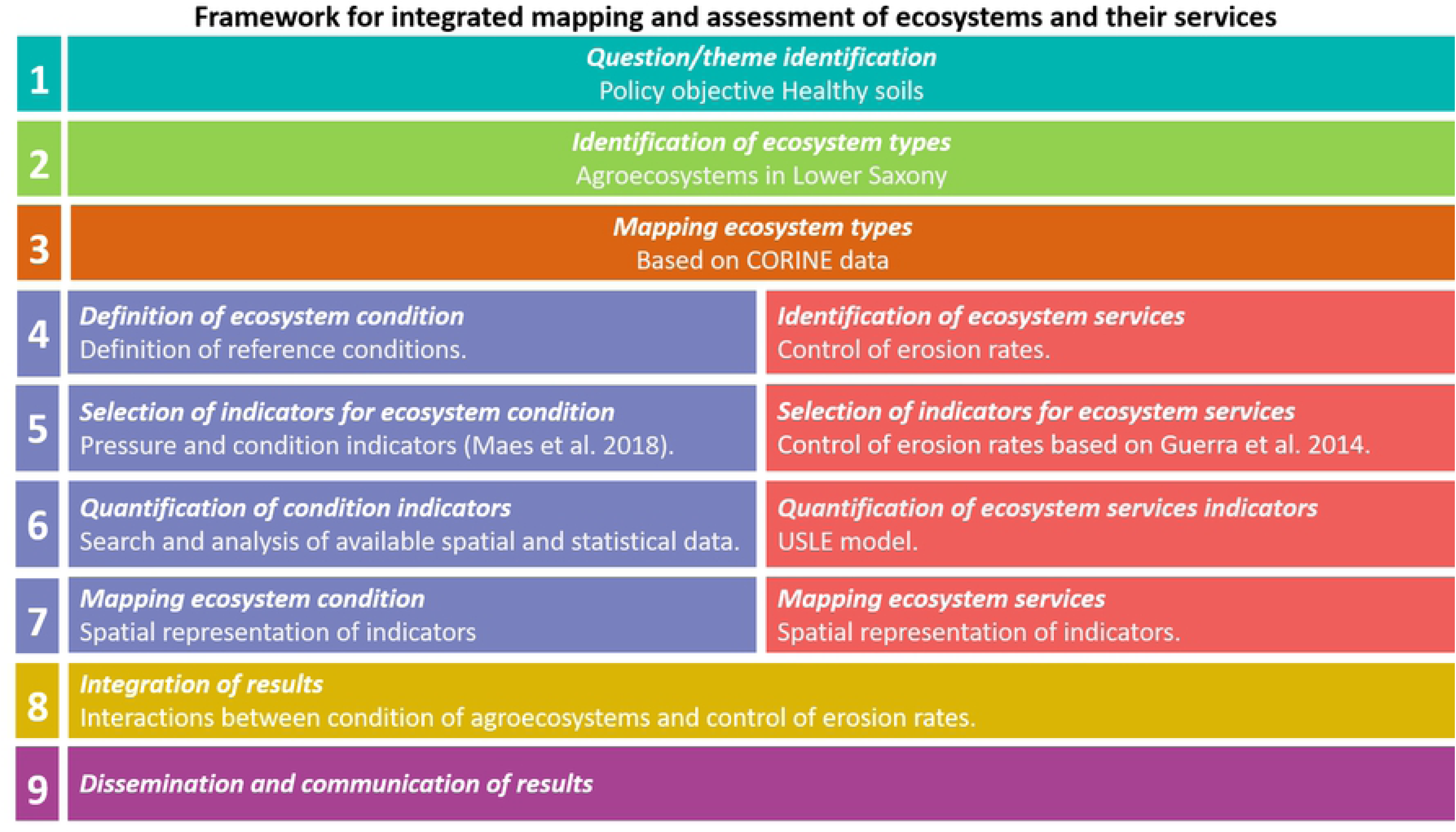
Conceptual framework applied for integrated mapping and assessment of agroecosystems and the ecosystem service *control of erosion rates* (based on Burkhard et al. [27]).

### 2.3. Theme identification: Policy Objective healthy soils (Step 1)

The first step of the operational framework refers to the *question and theme identification*, which must be addressed in the ecosystem assessment so that it is relevant for policy, society, business or science. In this case study, we identified the policy objective *maintaining healthy soils*. Healthy soils, especially for agriculture, have a high functionality, including high biodiversity, fertility and the capacity to sustainably deliver multiple ecosystem services. These services include food and fibre, climate and water regulation, water purification, carbon sequestration, nutrient cycling and provision of habitat for biodiversity [30]. At the same time, the delivery of other ecosystem services such as water supply and regulation, pollination and soil erosion regulation should not be impaired [31].

For this study, we focus on the ecosystem service *control of erosion rates.* The attributes of agroecosystem condition that affect the delivery of this ecosystem service depend on the condition of the soils and the presence of semi-natural areas within or in the vicinity of the agricultural fields [32, 33]. Additionally, the presence of livestock can affect this service by altering the structural condition of soils [34]. Other condition attributes that affect the provision of the service are certain crop rotations and crop types as well as the state of the landscape in which the agroecosystem is embedded [35]. In this case, the main typologies of environmental pressures, habitat and land conversion, climate change, input nutrients and pesticides, overexploitation, and introduction of invasive species, proposed by Maes et al [12] affect the condition attributes (see Fig 3).

**Fig 3.**
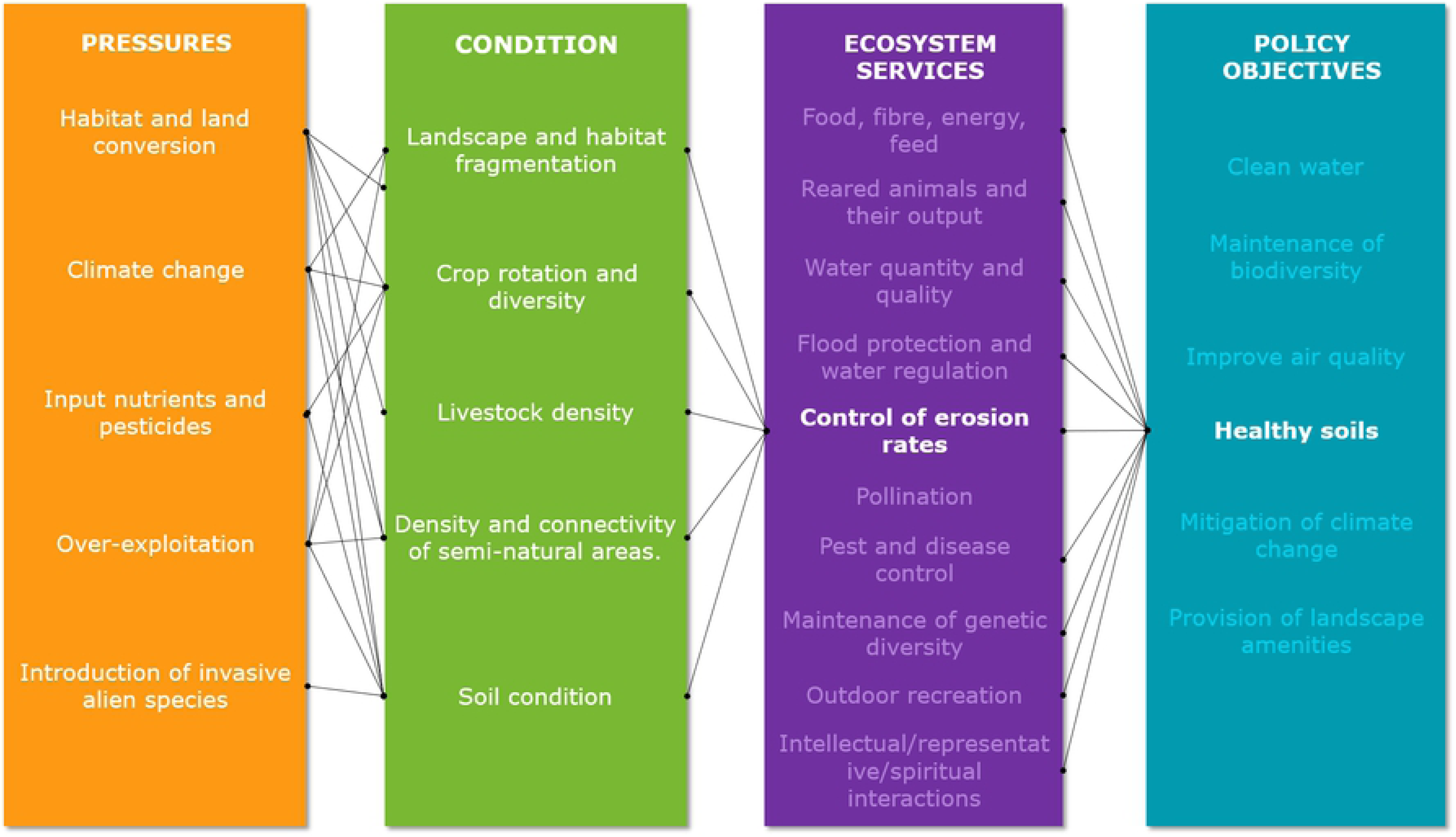
Synthesis of links between environmental pressures, condition, ecosystem service *control of erosion rates* and policy objectives in agroecosystems (based on Maes et al [12].

Healthy soils have been an important element in national and European policies. In Germany, for instance, the objective of maintaining and preserving healthy soils was established in the Federal Soil Protection Act of March 17^th^ 1998 [36] and the Federal Soil Protection and Contaminated Sites Ordinance of July 12^th^ 1999 [37]. Both regulations aim to sustainably secure and restore the soil functions by protecting soils against harmful changes and remediating contaminated sites. At EU level, the European Commission adopted the Thematic Strategy for Soil Protection [38] with the objective of protecting soils across the EU. Although the proposal for a Soil Framework Directive was withdrawn by the Commission in 2014, the 7^th^ Environmental Action Programme (EAP) came into force in 2014 [9]. The objective of the EAP is the protection and sustainable use of soils. Under priority objective 1, the EAP states that by 2020 “land is managed sustainably in the Union, soil is adequately protected and remediation of contaminated sites is well underway”. To do so, more efforts are required to reduce soil erosion and increase soil organic matter, as well as to remediate contaminated sites.

### 2.4. Identification and mapping of the ecosystem type agroecosystems (Steps 2 and 3)

The second step of the operational framework refers to the identification of the ecosystem type(s) to be assessed in this study. The ecosystem type is agroecosystems which are “communities of plants and animals interacting with their physical and chemical environments that have been modified by people to produce food, fibre, fuel, and other products for human consumption and processing” [39]. Maes et al.[11] proposed a classification of ecosystem types for MAES, in which cropland and grassland belong to the ecosystem type of agroecosystems. We selected cropland ecosystems in Lower Saxony as a study subject because croplands are a main provider of important ecosystem services such as biomass used as food, fodder or as energy source. Croplands are especially threatened by management and external pressures like droughts or floods related to climate change. Due to management induced bare soils within parts of the year, soils on cropland are especially affected by soil erosion.

The third step consists of the mapping of the ecosystem type agroecosystems that was previously identified. For this purpose, we used the CORINE Land Cover Data for the year 2012 [40], specifically the CORINE Land Cover type 2. Agricultural areas, which includes: 211. Non-irrigated arable land, 221. Vineyards, 231. Pastures, 242. Complex cultivation patterns, and 243. Land principally occupied by agriculture, with significant areas of natural vegetation (see Fig 1).

### 2.5. Definition of ecosystem condition and identification of ecosystem services delivered by agroecosystems (Step 4)

The fourth step of the operational framework refers to the definition of ecosystem condition and the ecosystem services delivered. Agroecosystems are usually purposely heavily modified ecosystems [18], which means that it is not feasible to compare their condition against undisturbed natural ecosystems. Table 1 shows the median values and the available reference values used in this study to determine the condition of the agroecosystems in Lower Saxony, based on the selected indicators.

**Table 1.** Indicators used for the assessment of environmental pressures and condition of agroecosystems and the ecosystem service *control of erosion rates* in Lower Saxony

| Indicator class | Indicator | Description | Units | Spatial resolution | Year | Median Lower Saxony | Reference value | Source |
| --- | --- | --- | --- | --- | --- | --- | --- | --- |
| Pressure indicators |  |  |  |  |  |  |  |  |
| Habitat conversion and degradation | Change in ecosystem extent | Change in the area (size) of the ecosystem within the years 2016 and 2017. | % per year | Municipality | 2016-2017 | -0.034 | N.A | [44] |
| Climate | Mean annual temperature | Annual mean of the monthly averaged mean daily air temperature in 2 m height above ground | °C | 1000 m | 1988-2018 | 9.74 | N.A | [45] |
|  | Mean annual precipitation | Annual sum of monthly precipitation. | mm |  | 1988-2018 | 765.7 | N.A |  |
|  | Drought index | Annual mean of drought index after de Martonne. | mm °C<sup>-1</sup> |  | 1995-2018 | 37.78 | Very humid (35-55) [46] |  |
|  | Precipitation 10 mm | Number of days with precipitation ≥ 10 mm per year. | Number of days |  | 1988-2018 | 19.45 | N.A |  |
|  | Precipitation 20 mm | Number of days with precipitation ≥ 20 mm per year. |  |  |  | 3.77 | N.A |  |
|  | Precipitation 30 mm | Number of days with precipitation ≥ 30 mm per year. |  |  |  | 0.91 | N.A |  |
|  | Begin of vegetation period | Number of consecutive days of the year. It indicates the begin of the first spring | Consecutive days of the year |  | 1992-2018 | 82.68 | N.A |  |
|  | Summer soil moisture | Modelled trend in soil moisture over a depth of 60 cm. | % NFK (usable field capacity) |  | 1991-2010 | 66.82 | N.A |  |
| Others | Soil erosion | Amount of soil loss per hectare in a year (Actual soil loss) | t ha<sup>-1</sup> per year | 50 m | 2010 | 0.13 | 0 - 3 [47] | [48] |
|  | Loss of organic matter | Percentage of soil organic carbon loss per year. | Mg C ha<sup>-1</sup> per year | 1000 m | 2014 | 0.015 | N.A | [49,50] |
| Ecosystem condition indicators |  |  |  |  |  |  |  |  |
| Structural ecosystem attributes (general) | Crop diversity | Average number of crops in a 10 km diameter per municipality | Number of crops | 50 m | 2018 | 44.41 | N.A | [51] |
|  | Density of semi-natural areas | Percentage of semi-natural areas | % | 50 m | 2012 | 13.16 | N.A | [40] |
|  | Share of fallow land in Utilized Agricultural Area (UAA) | Percentage of arable land that is not being used for agricultural purposes within the UAA. | % | Municipality | 2010 | 0.80 | N.A | [52] |
|  | Share of arable land in Utilized Agricultural Area (UAA) | Percentage of land used for the production of crops within the UAA | % | Municipality | 2010 | 81.32 | N.A | [52] |
|  | Share of permanent crops in Utilized Agricultural Area (UAA) | Percentage of land used for permanent crops within the UAA | % | Municipality | 2010 | 0 | N.A | [52] |
|  | Livestock Density | Stock of animals (cattle, sheep, goats, equidae, pigs, poultry and rabbits) converted in livestock units (LU) per hectare of UAA. | LU ha<sup>-1</sup> | Municipality | 2010 | 0.81 | N.A | [52] |
| Structural soil attributes | Soil Organic Carbon (SOC) | Concentration of topsoil organic carbon. | % or gr kg<sup>-1</sup> | 1000 m | 2010 | 2.03 | 1 - 2 % [53] | [54] |
|  | Soil erodibility | Susceptibility of soil to erosion by runoff and raindrop impact. | K factor [t h ha<sup>-1</sup> N<sup>-1</sup>] | 50 m | 2010 | 0.21 | N.A | [54,55] |
|  | Bulk density | Weight of soil per cubic meter | t m<sup>-3</sup> | 500 m | 2009 | 1.3 | 1.6 g cm<sup>-3</sup> for sandy and sandy loam soils<br>1.75 g cm<sup>-3</sup> for coarse textured soils (with clay content < 17.5%) [53] | [56,57] |
| Ecosystem services indicators |  |  |  |  |  |  |  |  |
| Control of erosion rates | Soil erosion risk | Potential soil loss. | t ha<sup>-1</sup> per year | 50 m | 2010 | 0.81 | N.A | [48] |
|  | Prevented soil erosion | Difference of potential and actual soil loss | t ha<sup>-1</sup> per year | 50 m | 2010 | 0.67 | N.A | [54] |
|  | Provision capacity | Share of mitigation of soil erosion (0 to 1) | Dimensionless | 50 m | 2010 | 0.85 | From 0 (no mitigation) to 1 (complete mitigation) | [54] |

We selected the ecosystem service *control of erosion rates,* because soil erosion is one of the main threats to soils [38] with negative impacts on crop production, water quality, mudslides, eutrophication, biodiversity and carbon stock loss [26]. Soils are the medium on which crops are grown and their functionality is the base for biomass production, storage, filtration and transformation of nutrients and water. Furthermore, healthy soils are important for biodiversity conservation and act as carbon storage pools. Additionally, soils are the platform for human activities, provide raw materials and store geological and archaeological heritage [41]. Therefore, soil degradation will lead to the declines of many ecosystem services [42]. Soil erosion causes the soil surface to decrease and then reduce its thickness. Especially the loss of the humus-rich, fertile topsoil layers leads to a reduction of soil functionality and the capacity to provide ecosystem services. If soil formation is not able to compensate soil losses, soil erosion will threaten sustainable crop production as well as water regulation and filtration capacities [43].

### 2.6. Selection of indicators for agroecosystem condition and the ecosystem service control of erosion rates (Step 5)

The fifth step of the operational framework refers to the selection of the indicators for the assessment of ecosystem condition and ecosystem services. As one of the aims of this study is to test the framework and indicators proposed by MAES for mapping and assessment of ecosystem condition in the EU, we chose the indicators for pressures and condition of agroecosystems presented in the 5th MAES report [12]. We selected the indicators for the ecosystem service *control of erosion rates* based on existing literature and the frameworks used by Guerra et al. [58] and Steinhoff-Knopp and Burkhard [25].

#### 2.6.1. Criteria for selecting indicators

Ecosystem condition indicators allow to assess the overall quality of an ecosystem and its main characteristics that underpin its capacity to deliver ecosystem services [1]. These indicators, together with information on ecosystem extent and services, constitute the main inputs for integrated ecosystem assessments that analyse the links between ecosystem condition, habitat quality and biodiversity, ecosystem services and the consequences for human wellbeing [27].

Maes et al. [12] highlighted the main characteristics that ecosystem condition indicators must have in order to inform policies related to the use and protection of natural resources. First, ecosystem condition indicators need to be aligned with the MAES conceptual framework in which socioeconomic systems are linked with ecosystems through the flow of ecosystem services and the drivers that affect ecosystems. Second, they need to support the objectives of the EU environmental legislation and the objectives of the natural capital accounts. Third, they need to be policy-relevant, which means that they support EU environmental policies as well as related national policies and other policies. Fourth, they need to be spatially explicit by considering the distribution of ecosystems and their use, and they need to be specific for each ecosystem type. Fifth, they need to contribute to measuring progress/trends against a policy baseline towards different policy targets.

For this research purpose which aims to assess the feasibility of the MAES indicators proposed by Maes et al [12] to estimate the condition of agroecosystems in Lower Saxony and to link them specifically with the ecosystem service *control of erosion rates*, we adopted the following criteria for selecting the indicators to be used in this case study:

- Relevancy: Indicators must be relevant to the ecosystem service *control of erosion rates*. This means that there is a clear connection between the condition parameter and the provision of the ecosystem service. This connection was determined based on the authors’ knowledge and evidence in scientific literature.
- Availability of data at a regional scale: The used data have an appropriate resolution for the study area and are detailed enough to recognize regional features (i.e. spatially explicit or data at the level of municipalities).
- Quantifiable: Indicators must be quantifiable and data can be compared among municipalities.
- Reliability: The quantification and monitoring of the indicators are both reliable (i.e. data obtained from officially reported data sets).

A total 23 indicators was selected for the assessment, whereof 11 indicators correspond to pressure indicators (including 7 climate indicators), 9 correspond to ecosystem condition indicators and 3 to the ecosystem service *control of erosion rates*. Table 1 provides detailed descriptions of the indicators, including the spatial resolution, year of data collection and data sources.

### 2.7. Quantification of indicators (Step 6)

The sixth step of the operational framework refers to the quantification of indicators for condition and the ecosystem service *control of erosion rates*. For this purpose, we conducted a data search in multiple public databases of the EU, Germany and Lower Saxony. In this section, we describe the calculation of each indicator.

#### 2.7.1. Environmental pressure indicators

*Change in ecosystem extent* was calculated based on the agricultural area data of the municipalities in Lower Saxony for the years 2016 and 2017. These data were obtained from the State Statistical Office of Lower Saxony [44].

### Climate

The data of the climate indicators were obtained from the German Weather Service (Deutscher Wetterdienst DWD) [45]. The detailed description of the indicators is provided below (see Table 1 for information on spatial resolution):

*Mean annual temperature:* Corresponds to the annual mean of the monthly averaged mean daily air temperature in 2 m height above ground given in degree Celsius, for the period between 1988 and 2018.

*Mean annual precipitation:* Corresponds to the annual sum of monthly total precipitation given in mm for the period between 1988 and 2018.

*Drought index:* The data of annual drought index (from de Martonne [46]) dMI was
calculated with:

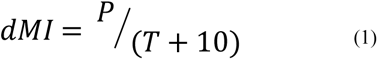

Where T [in degree Celsius] was obtained from temperature grids and P [in mm] from precipitation grids for the period between 1995 and 2018.

*Precipitation 10 mm, 20 mm and 30 mm:* Corresponds to the number of days with precipitations equal or higher than 10, 20 and 30 mm respectively, averaged for the period between 1988 and 2018.

*Begin of vegetation period:* Corresponds to the consecutive days of the year in which the first spring begins averaged for the period between 1992 and 2018.

*Summer soil moisture* was obtained from the DWD who used the AMBAV model that calculates the evapotranspiration and the soil-water balance for different crops [59]. This was done based on the soil moisture in 60 cm depth under grass derived from selected stations for the period between 1991 and 2010, for the months of June, July and August.

*Soil erosion* corresponds to the mean actual soil loss and was calculated with the Universal Soil Loss Equation (USLE), applying the German standard DIN 19708 [60].

The loss rates were modelled as raster GIS layers for Lower Saxony in a resolution of 50 m based on the methodology applied by [48]. The USLE was calculated as:

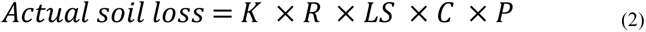

Where

*K* is the soil erodibility factor [t h ha^-1^ N^-1^] for Germany based on the soil overview maps (Bodenkundliche Übersichtskarten [61]) and the approach of Auerswald & Elhaus [55]

*R* is the rain erosivity factor [N ha^-1^ per year] for Germany based on mean annual summer precipitations for 1980-2010 from the DWD and the regression equation from standard DIN 19708.

*LS* is the topography factor (length and slope) [dimensionless] based on a 50 m Digital Elevation Model (DEM) and the approach from the DIN 19708.

*C* is the crop management factor [dimensionless] based on crop rotations from culture portions on the level of the municipality for the year 2010.

*P* is the erosion control practices factor [dimensionless] considered in this study as 1 due to lack of detailed data for the resolution we used.

The results from the calculations were clipped to the arable land according to the land use type *Non-irrigated arable land (2.1.1)* of the CORINE Land Cover 2012 [40].

*Loss of organic matter:* This indicator was obtained from the average eroded soil organic carbon dataset calculated for the EU [49]. This was done by applying the CENTURY model, which simulates the carbon and nitrogen dynamics in cultivated or natural systems, coupled with soil erosion [50]. We calculated the average soil organic carbon loss per municipality based on a 1 km raster data set.

#### 2.7.2. Ecosystem condition indicators

*Crop diversity* was calculated based on the number of field blocks in the year 2018. These data were obtained from the Land Development and Agricultural Support programme of Lower Saxony (LEA) Portal [51] of the Lower Saxonian Ministry of Food, Agriculture and Consumer Protection. We calculated this indicator by estimating the number of crops in a regular 1km raster by applying a moving window analysis with a radius of 5 km. By counting the overlaying field, we added them up and obtained the diversity of crops in a diameter of 10 km around the midpoints of each raster cell. The raster data were combined with the shapes of the municipalities to calculate the average number of crops per municipality.

*Density of semi-natural areas* was calculated based on the CORINE Land Cover data of 2012. The land cover classes were selected according to those proposed by García-Feced et al [62]: 2.3.1. Pastures, 2.4.3. Land principally occupied by agriculture, with significant areas of natural vegetation and natural grasslands, the land cover class 2.4.4. Agro-forestry areas is not present in the dataset in Lower Saxony. The data of these land cover classes were combined with the shapes of the municipalities to calculate the percentage of semi-natural areas in relation to the area of the municipality.

The *share of fallow land in Utilized Agricultural Area (UAA)* was calculated by dividing the number of hectares of fallow land and set aside or decommissioning land over the number of hectares of UAA per municipality. The same approach was used for the calculation of the share of arable land and permanent crops in the UAA. The data for the estimation of these indicators was obtained from the Thünen-Atlas (Collection of agricultural data from Germany) [52, 63] for the year 2010.

*Livestock density* was calculated by dividing the number of livestock units over the number of hectares of UAA per municipality, both were obtained from the Thünen-Atlas [52, 63] for the year 2010.

*Soil Organic Carbon* was calculated based on the datasets on *humus content in the topsoil* and the *usage-differentiated soil survey map* obtained from the Federal Institute for Geosciences and Natural Resources (BGR for its acronym in German) [54]. We used depths from 0 to 10 cm below the soil surface for grassland, pasture and forestry and depths from 0 to 30 cm under the soil surface for crop lands. In order to convert the humus content data into soil organic carbon data, we used the following equations:

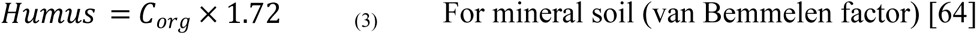

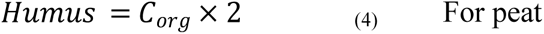

*Soil erodibility* was calculated as the K factor explained before in the USLE equation [48].

*Bulk density* was obtained from the bulk density calculated for the EU [57]. This physical property was derived from soil texture data sets as described by Ballabio et al [56].

#### 2.7.3. Ecosystem service indicators

The ecosystem service *control of erosion rates* is a regulating ecosystem service that mitigates the structural impact *potential soil loss.* In this study, we adapted the conceptual framework for assessing the provision of regulating ecosystem services developed by Guerra et al [58]. Fig 4 is a graphical representation of the adapted framework, where A shows the concept for the assessment of the ecosystem service *control of erosion rates* and B shows the implementation in this study.

**Fig 4.**
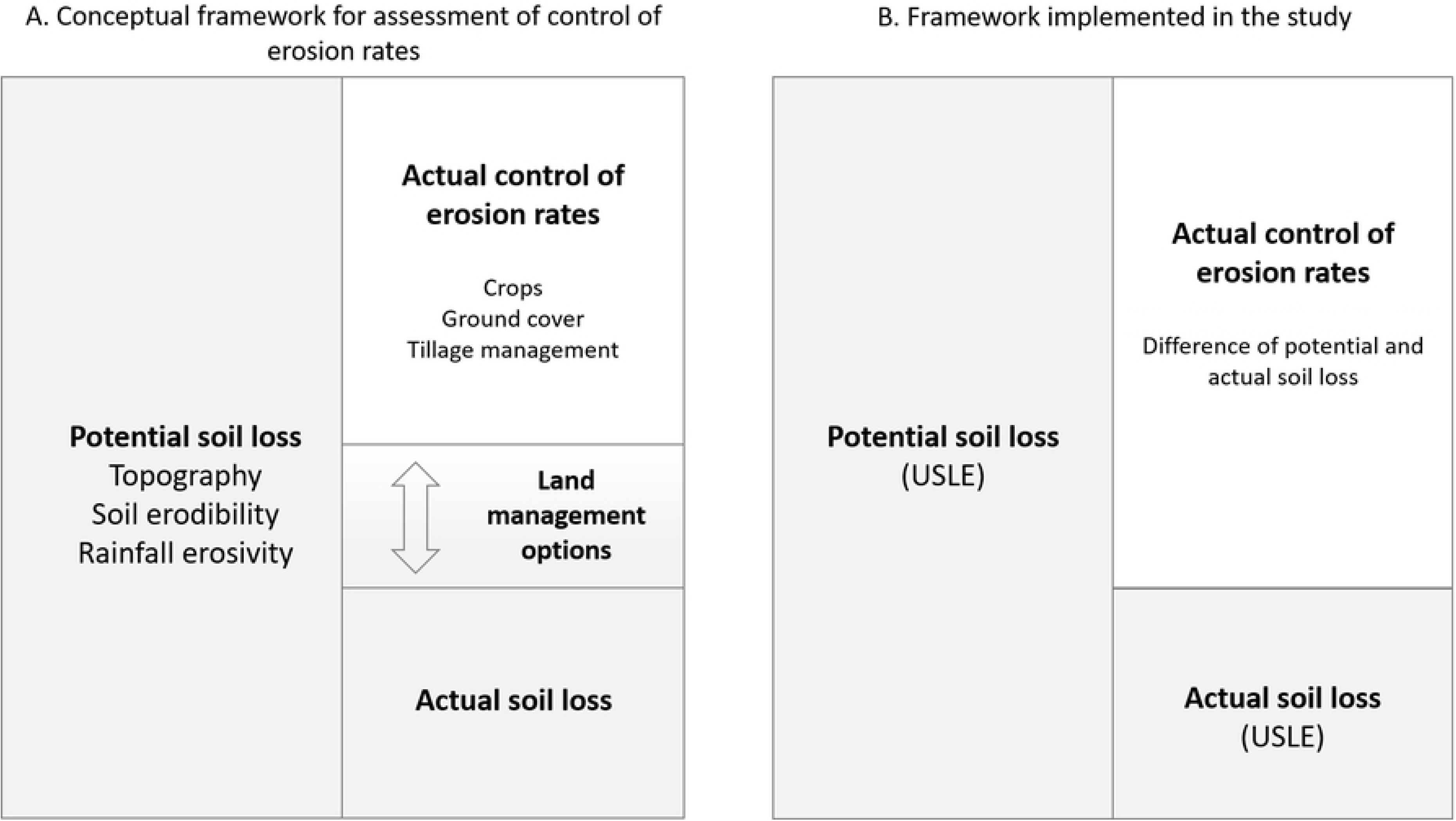
Conceptual framework applied for the assessment of the ecosystem service *control of erosion rates* based on Guerra et al.[58].

*Soil erosion risk* is the mean potential soil loss defined as the amount of soil that is lost when there is no vegetation covering the ground [25]. Potential soil loss is determined by the USLE factors rain erosivity (R), soil erodibility (K) and topography (LS) as shown in the following equation [48] (also described in section 2.7.1 for the pressure indicator *soil erosion*):

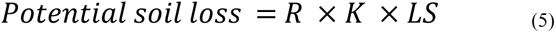

The results from the calculations were clipped to the arable land according to the land use type *Non-irrigated arable land (2.1.1)* of the CORINE Land Cover 2012.

The *actual control of erosion rates* denotes the prevented soil erosion calculated as the difference of *potential soil loss* (soil erosion risk) and *actual soil loss* (soil erosion - described in section 2.7.1).

Another indicator used to calculate the ecosystem service *control of erosion rates* is the *provision capacity.* This is defined as the fraction of the potential soil loss that was mitigated by the actual service provision. It ranges from 0 to 1, where 0 represents no mitigation and 1 complete mitigation and was calculated based on the USLE model results for Lower Saxony (see [25] for more detailed explanation of the approach).

The data of these indicators were combined with the shapes of the municipalities to calculate the average value per municipality.

### 2.8. Mapping of indicators (Step 7)

This step of the operational MAES framework refers to the spatial visualisation of the ecosystem condition and the ecosystem service *control of erosion rates* indicators in maps. All the indicators were edited in ArcGIS 10.7 for representation and analysis and were aggregated to the level of the municipalities to facilitate a comparison.

### 2.9. Integration of results (Step 8)

The integration of results refers to the analysis of relationships and interactions between the condition of agroecosystems and the supply of the ecosystem service *control of erosion rates*. This analysis was conducted from two different angles: one assesses the statistical correlations, the other one analyses the spatial distributions and relationships from the compiled maps.

For the statistical correlations, seven classes of the ecosystem service *control of erosion rates* were selected, based on the classification proposed by Steinhoff-Knopp and Burkhard [25] for the potential soil erosion by water on cropland in Lower Saxony. The classes are *No to very low* (less than 1 t ha^-1^ per year eroded soil), *very low* (1 to 5 t ha^-1^ per year), *low* (>5 to 10 t ha^-1^ per year), *medium* (>10 to 15 t ha^-1^ per year), *high* (>15 to 30 t ha^-1^ per year), *very high* (>30 to 55 t ha^-1^ per year) and *extremely high* (≥ 55 t ha^-1^ per year). We considered the environmental pressures and ecosystem condition indicators per class, which are also used to classify the actual *control of erosion rates*. We performed the Akaike Information Criterion

(AIC) [65] technique to estimate the likelihood of the different pressure and condition indicators to predict the values of the seven *control of erosion rates* classes mentioned above. Additionally, we performed the Kruskal-Wallis rank test [66] on the class medians to detect significant differences between the seven classes and the Jonckheere-Terpstra test [67, 68] to identify positive or negative relationships between the delivery of the ecosystem service *control of erosion rates* and the environmental pressures and ecosystem condition. All statistical work was conducted in RStudio (version 1.2.1335) [69].

In order to show an integrated overview of the distribution of the ecosystem service *control of erosion rates*, the potential soil loss, and the pressures and condition in Lower Saxony, we normalized all the variables and standardized them to a 0 to 100 scale to make them comparable. When looking at pressures and condition variables, we considered the presumable effect of each of them based on the supply of the ecosystem service *control of erosion rates* to do the normalization. The pressure indicators *drought index* and *summer soil moisture*, for instance, are thought to have a positive effect on the supply of the ecosystem service. Regarding the drought index (Ia DM) in regions classified as arid (Ia DM<10) and semi-arid (10≤ Ia DM<20) in the *index of the Martonne*, the protective cover provided by plants against rain splash decreases with increased aridity [70]. This means that the higher the value of the drought index, the lower the erosion risk. Similarly, higher soil moisture maximizes vegetation cover, resulting in the minimization of sediment transport capacity, with important differences between clay and sandy soil textures [71]. On the other hand, the condition indicators *fallow land*, *livestock density* and *soil erodibility (K factor)* have a presumably negative effect on the *control of erosion rates.* Poor structural stability as well as less plant cover in fallow systems result in an increased erosion risk [72], and a higher percentage of fallow land, indicating areas with bare soil, increases the soil erosion in a specific area. Likewise, the trampling of livestock disturbs soil and loosens it, which makes the soil easier to remove by agents of transport and therefore increases its erodibility [34]. These five indicators were multiplied by -1 in order to take into account the positive or negative effects when normalizing the original data. We then calculated the average of the normalized values for the pressure and condition indicators.

In order to spatially visualize the relationships between the indicators, we created maps showing the overlaps of pressures, condition, soil erosion risk and provision capacity. This analysis allowed us to identify how pressures and condition were related to soil erosion risk and the provision capacity of the agroecosystems to control soil erosion.

### 2.10. Dissemination and communication of results (Step 9)

This step refers to the preparation of maps and other accompanying material for an effective dissemination and communication of the results. According to Burkhard et al.[27] the results must be communicated to decision makers and other stakeholders who might be interested in the them in order to answer the initial question(s) posed in Step 1. However, for this assessment, we have not involved stakeholders and these results have not been communicated.

## 3. Results

The results of the assessment are presented in maps showing the distribution of the indicators of environmental pressures, ecosystem condition and the ecosystem service *control of erosion rates* within Lower Saxony. Other graphs and maps show the integration of results and the relationships between pressures, condition and *control of erosion rates* (see data per municipality in Supplementary material S1).

### 3.1. Mapping and assessment of agroecosystem condition in Lower Saxony

#### 3.1.1. Pressure indicators

##### Change in ecosystem extent

Agroecosystems in Lower Saxony did not experience great changes in extent from the year 2016 to 2017. Most of the municipalities showed changes in agroecosystem extent from 0% to +/- 0.6%. The changes were more significant in municipalities such as Solling (district of Northeim), in the south, with an increase from 199 ha to 291 ha (+46%); Rehlingen (district of Lüneburg), in the northeast, from 1441 ha to 2085 ha (+45%); and Osterheide (district of Heidekreis), also in the northeast with an increase from 754 ha to 1000 ha (+33%). On the other hand, there were significant reductions in the size of agroecosystems on the North Sea islands of Baltrum from 134 ha to 56 ha (−58%) and Juist from 1232 ha to 273 ha (- 77%), both in the district of Aurich (Fig 5a).

**Fig 5.**
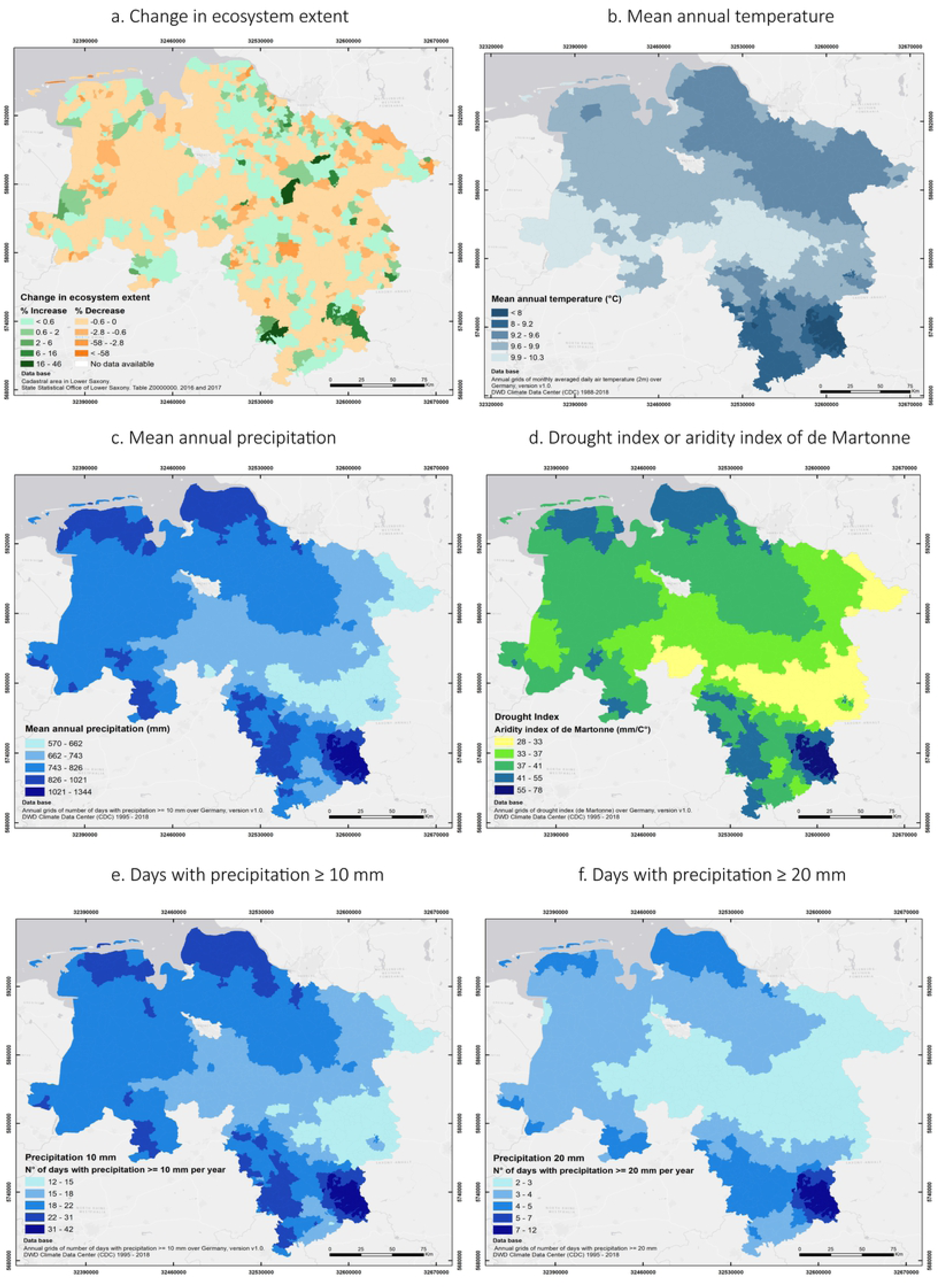
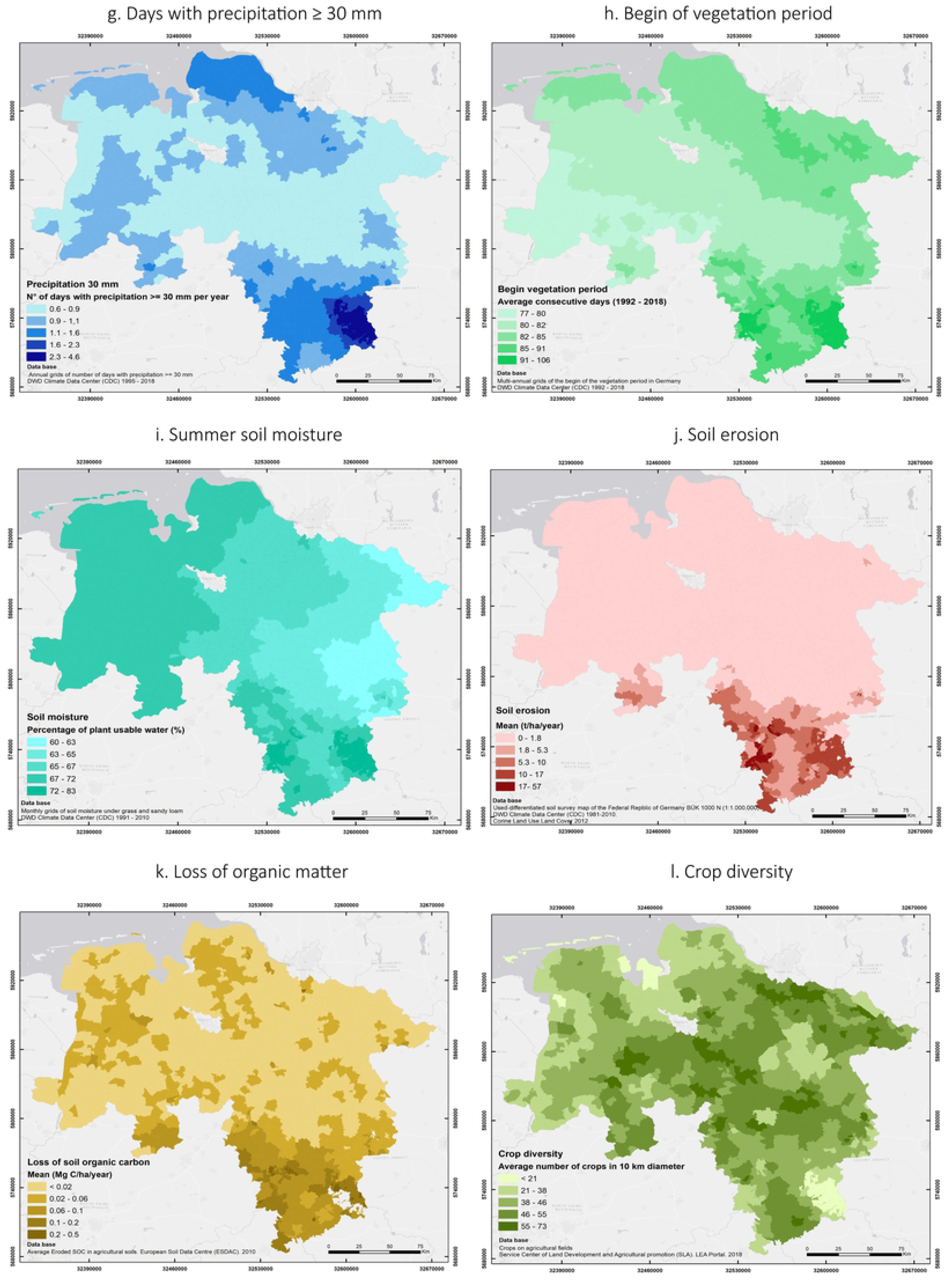
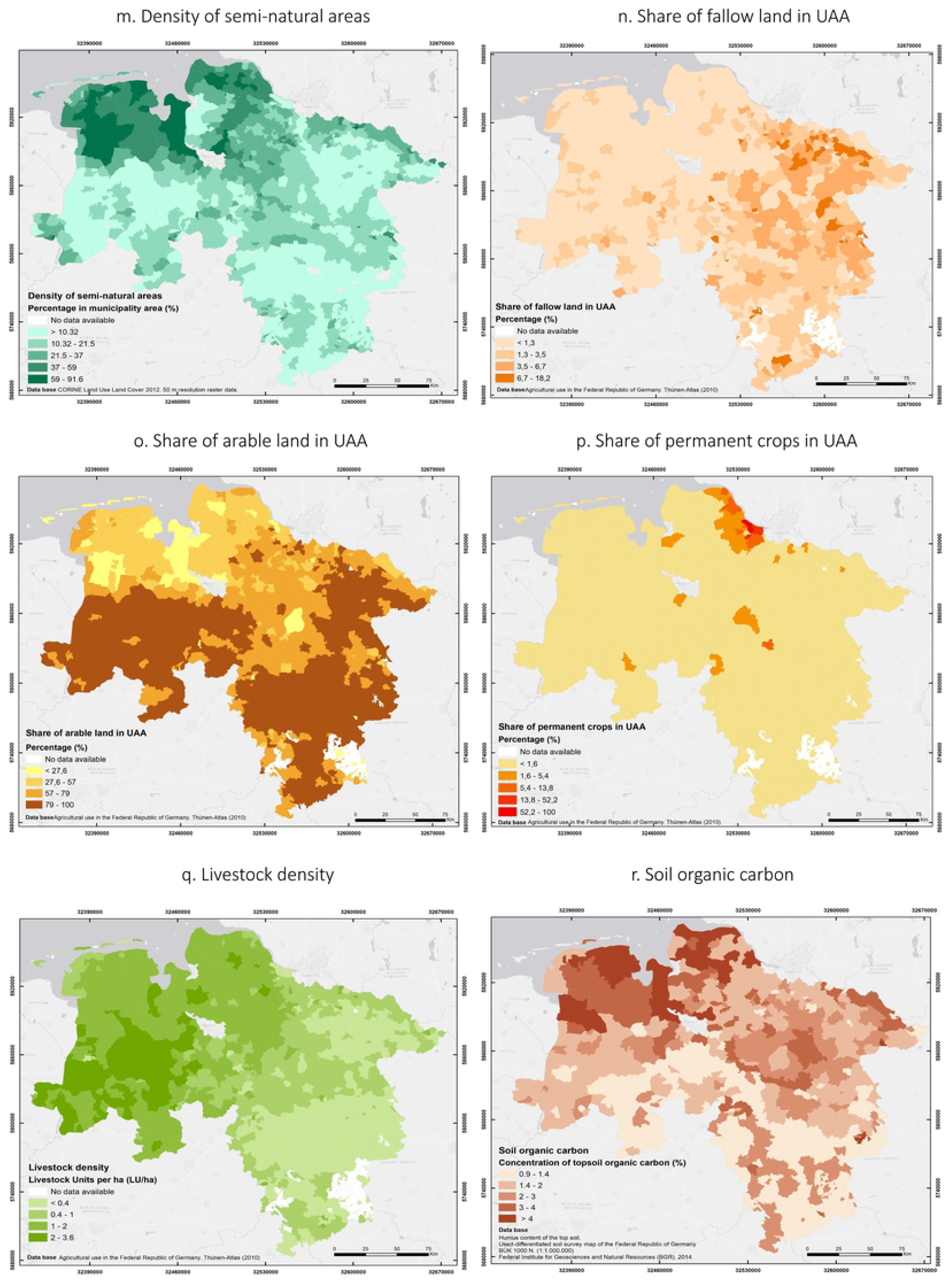
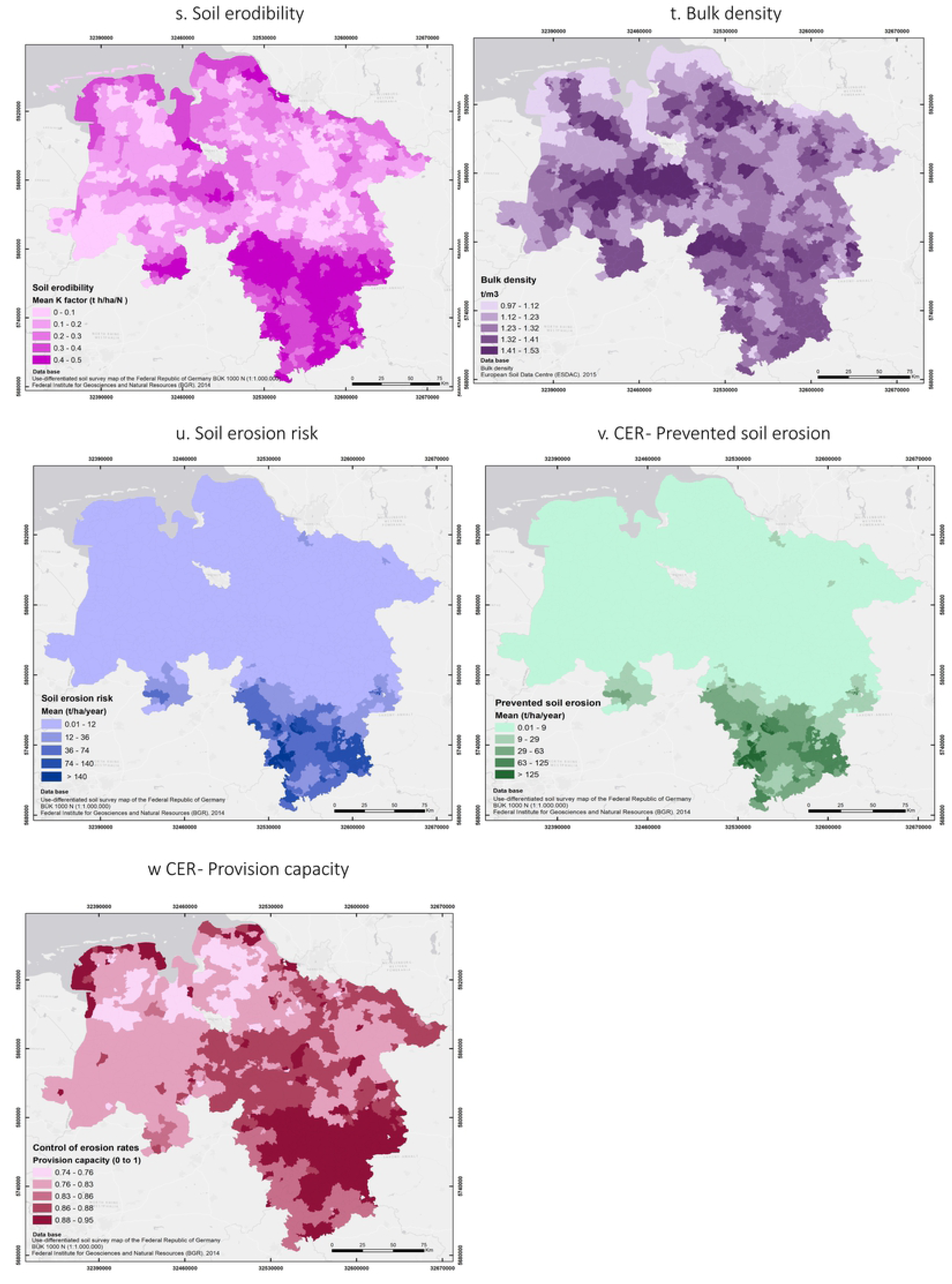
Maps of indicators of environmental pressure, ecosystem condition and control of erosion rates in Lower Saxony. (Large maps in Supplementary material S2)

##### Climate

*Mean annual temperatures* in Lower Saxony ranged from 6.7 °C to 10.3 °C, with the lowest temperatures recorded in mountainous areas such as the Harz (district of Goslar) in the southeastern part (Fig 5b).

*Mean annual precipitation* in Lower Saxony ranged from 570 mm in the east to 1344 mm in the mountainous region located in the southeastern part. Precipitation in the coastal areas and mountainous regions in the south ranged from 826 mm to 1021 mm (Fig 5c).

*Drought index or aridity index of the Martonne* ranged from humid (28) to extremely humid (78) across the study area. Municipalities located in mountainous regions (Braunlage and Clausthal-Zellerfeld in the district of Goslar) had the lowest aridity together with areas with continental weather (near the state Saxony Anhalt and some areas close to North Rhine Westphalia) (Fig 5d).

The number of days with precipitation higher than 10, 20 and 30 mm was negatively correlated with the amount of rainfall that was recorded. For instance, the number of days with precipitation between 10 and 20 mm ranged from 2 to 42 days, while the number of days with precipitation between 20 mm and 30 mm ranged from 0.6 to 4.6 days (Fig 5e-g).

Similarly to other climatic parameters, higher precipitations occurred in mountainous and coastal regions.

The *begin of the vegetation period* in Lower Saxony ranged from 77 to 106 consecutive days of the year in the period between 1992 and 2018 across the different municipalities. The areas with the latest beginning of spring are located in the mountainous regions (Braunlage and Clausthal-Zellerfeld in the district of Goslar), which is related to the prevalence of lower temperatures throughout the year (Fig 5h).

*Summer soil moisture* in Lower Saxony was in line with the precipitation levels showing percentages of plant-available water ranging from 60% in the eastern part to 83% in the southeast (mainly mountainous regions) (Fig 5i).

*Soil erosion* had on average the highest values in the southern mountainous region (56.9 t ha^-1^ per year in the municipality of Wenzen, district of Holzminden), whereas municipalities located in the Lower Saxonian German Plain in the northern half of the state, showed median values below 1.8 t ha^-1^ per year of (Fig 5j).

*Loss of organic matter* had the highest values in the southern region (0.48 Mg C ha^-1^ per year, municipality of Braunlage, district of Goslar). Other municipalities in the east and north showed values ranging from 0.06 to 0.1 Mg C ha^-1^ per year. The median for the whole federal state was 0.02 Mg C ha^-1^ per year (Fig 5k).

#### 3.1.2. Ecosystem condition indicators

*Crop diversity* varied greatly across Lower Saxony. The highest values are located in the northeastern and central areas ranging from 55 to 73 crop species. Whereas the lowest values, below 21 crop species, were recorded on the islands in the northwest and the mountainous area in the district of Goslar in the southeast (Fig 5l).

*Density of semi-natural areas* was higher in the northwest of Lower Saxony, with percentages as high as 91% in the municipality of Ovelgönne (district of Wesermarsch) and more than 87% in Engelschoff (district of Stade), mostly represented in the form of pastures. Almost 40% of the municipalities distributed in the southern and northeastern part have less than 10% of semi-natural areas and feature intensive agriculture, mainly with cereal crops (Fig 5m).

The *share of fallow land in UAA* was relatively low in Lower Saxony, with a median of 0.86% for the whole federal state. Almost 60% of the municipalities had a percentage of fallow land below 1.3%, whereas only 41 municipalities had values higher than 6.7%, mainly located in the eastern region (Fig 5n).

The *share of arable land in UAA* was higher in areas of lower density of semi-natural elements, which is in line with theoretical expectations, where most regions with intensive agriculture tend to have low values of semi-natural vegetation [62]. Municipalities with a share of arable land of around 80% were mainly distributed in the east and southwest parts of the federal state, where the production of crops such as grains and wheat was high (Fig 5o).

*Share of permanent crops in UAA* was low in Lower Saxony, with only five municipalities located in the district Stade in the northern part, with a share higher than 80% that correspond mainly to crops such as tree and berry fruits and fruit tree plantations in the fruit growing region “Altes land”. The median share of permanent crops for the federal state was 0%, since only 175 municipalities out of the 959 in the study have values higher than 0% (Fig 5p).

*Livestock density* was higher in the western part of Lower Saxony with the highest number of livestock units per hectare (LU ha^-1^) in the districts of Vechta and Cloppenburg with values between 2 and 3.6 LU ha^-1^. Values smaller than 0.4 LU ha^-1^ were recorded in the eastern part of the state and on some islands in the northwest. The median livestock density of the federal state was 0.8 LU ha^-1^ (Fig 5q).

*Soil Organic Carbon* concentrations were high in the northwest of the state with levels of topsoil organic carbon higher than 4% (Fig 5r).This agrees with large peatland areas in Northwestern Germany. The levels were higher than the threshold values between 1 and 2% as estimated by Kibblewhite et al [53]. On the other hand, levels of soil organic carbon in the eastern part of the study area, especially in the district of Wolfenbüttel, were lower than 0.9%, which could be an indication of potential degradation.

*Soil erodibility* or K factor values ranged from 0 to 0.5 t h ha^-1^ N^-1^ and showed higher mean values in the southeast of the study area and some municipalities in the northwest (Fig 5s). Additionally, the topsoils in these areas had high contents of silt, which makes them highly erodible [24].

*Bulk density* in Lower Saxony ranged from 0.97 t m^-3^ to 1.53 t m^-3^ distributed along all the municipalities (Fig 5t). These values were lower than the threshold levels for sandy and sandy loam soils of 1.6 g cm^-3^ estimated by Huber et al [47].

#### 3.1.3. Ecosystem service indicators

*Soil erosion risk* showed high values in the southeast part of the study area with values higher than 140 t ha^-1^ per year, which are considered extremely high according to Steinhoff- Knopp & Burkhard [25]. However, the median value of the entire state was calculated to be 0.8 t ha^-1^per year, and some municipalities even showed mean values as low as 0.01 t ha^-1^ per year (Fig 5u).

*Prevented soil erosion* showed a concentration of high values mainly in the southeastern part of the study area (Fig 5v). As previously mentioned, this area also showed high values of potential and actual soil losses. However, the actual soil loss was considerably lower than the calculated soil loss potential, resulting in a high ecosystem service provision in this area.

*Provision capacity* was relatively high in Lower Saxony with values ranging from 0.74 to 0.95 across all municipalities (Fig 5w). The mean value of 0.85 indicates that most parts of the study area are effectively protected against soil erosion.

### 3.2. Relationships between agroecosystem condition and control of erosion rates

#### 3.2.1. Analysis of the relationships between indicators

In order to understand the relationships between agroecosystem condition and supply of the ecosystem service *control of erosion rates*, we analysed the likelihood of the different indicators (excluding soil erosion, soil erosion risk, and soil erodibility which were partly included in the calculation of the ecosystem service) to predict the classes from *no to very low* to *extremely high* mentioned in section 2.9. Our results show that the indicators that best predicted these classes are: *loss of organic carbon*, followed by *mean annual temperature* and *begin of vegetation period*. In contrast, the indicators that had the lowest likelihood to predict the classes were *mean annual precipitation* and *crop diversity* (see AIC rank on Fig 6 and Supplementary material S3).

**Fig 6.**
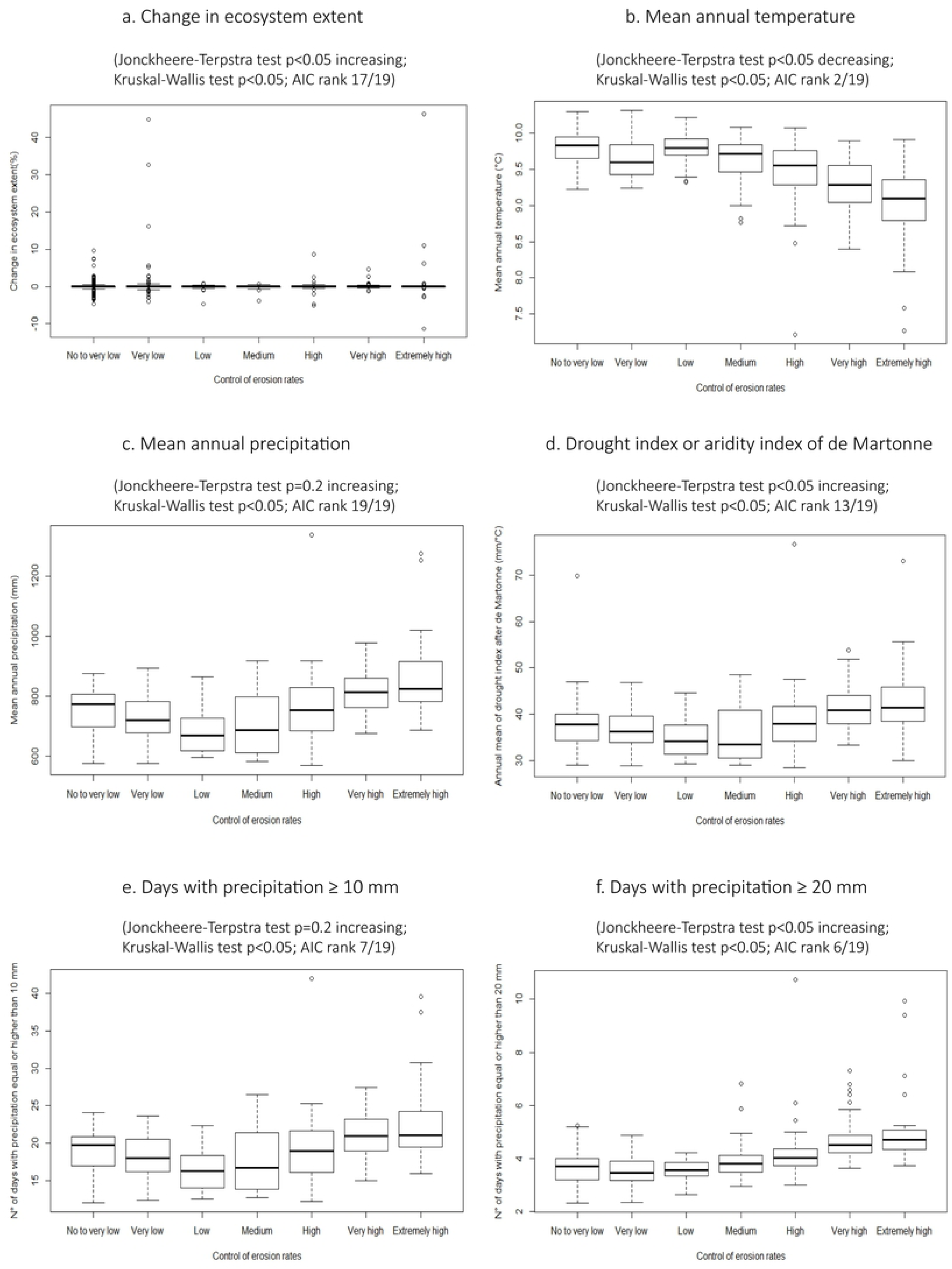
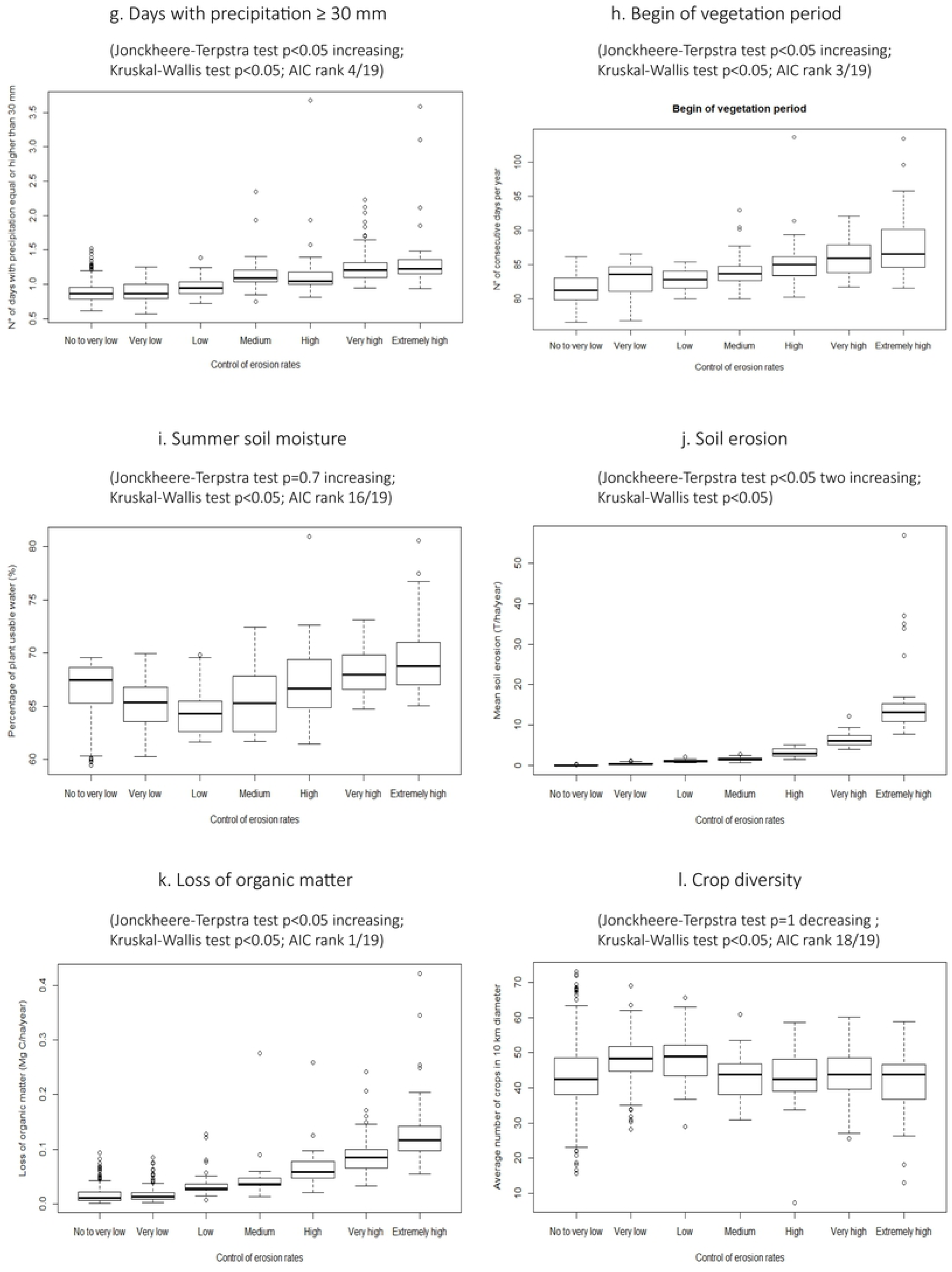
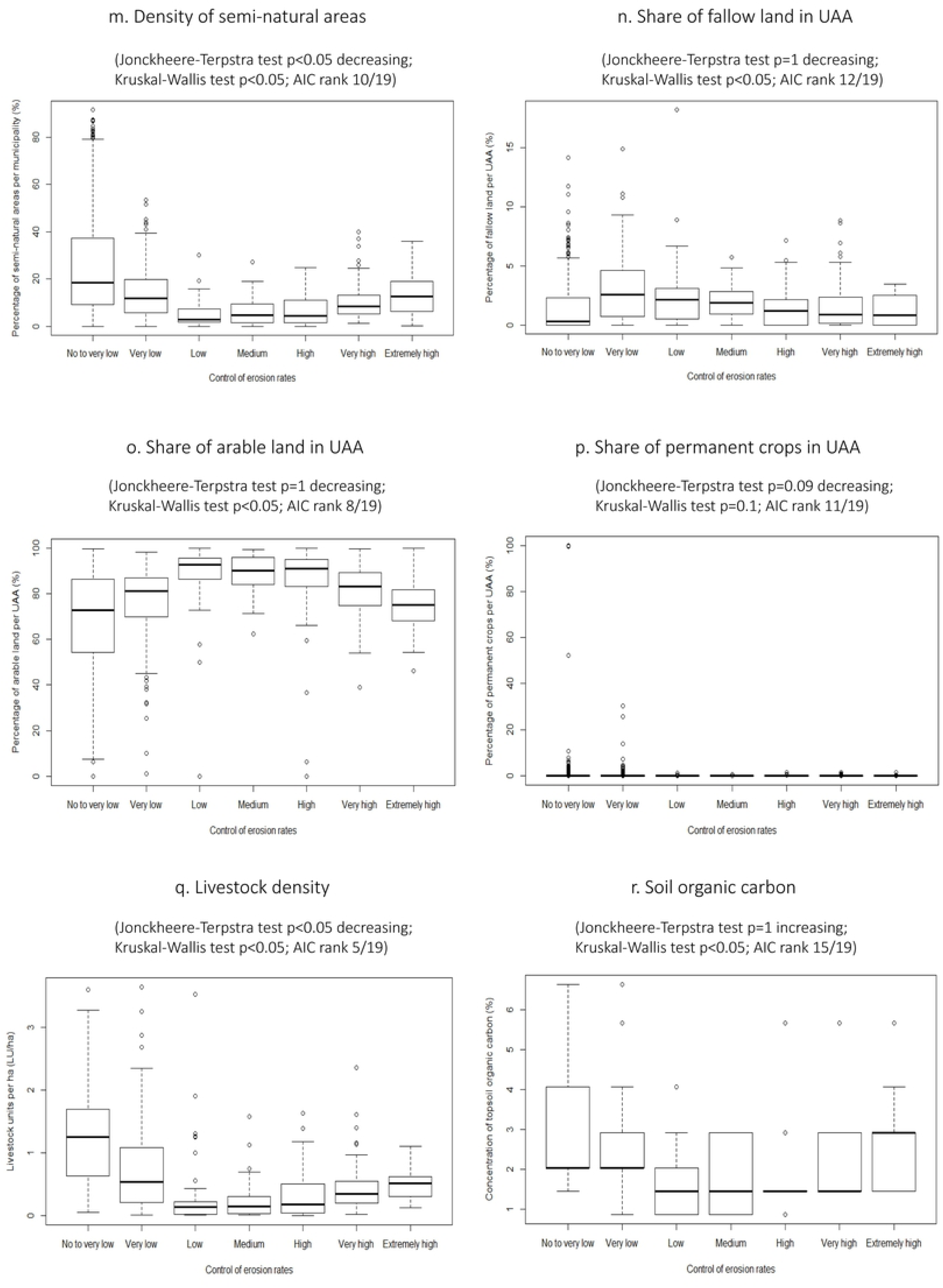
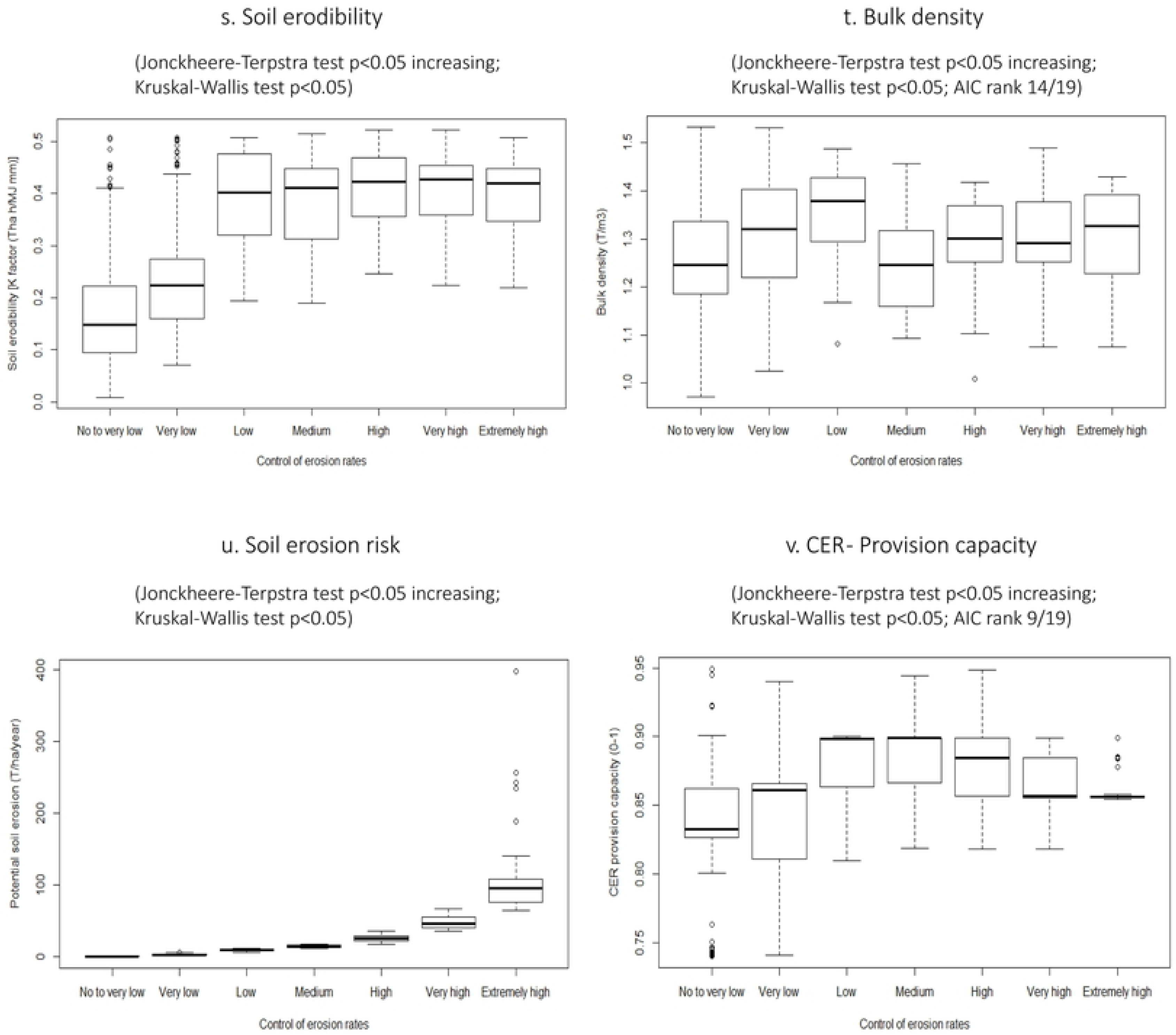
Relationships between the indicators of environmental pressures and condition, and the ecosystem service control of erosion rates.

Additionally, we analysed the relationships between pressures and ecosystem condition and the ecosystem service *control of erosion rates* (see Supplementary material S3). Our results show that the rates of control of erosion were slightly higher in areas with increased *ecosystem extent* (p < 0.05) (Fig 6a). For the climatic variables, negative, positive as well as not significant correlations were evident. For instance, in our study area, the *control of erosion rates* was extremely high in areas with lower temperatures (p < 0.05) (Fig 6b). On the other hand, high and extremely high *control of erosion rates* occurred in areas where variables such as *drought index*, *days with precipitations higher than 20 and 30 mm* and *begin of vegetation period* were high (p < 0.05). However, the relationships between *mean annual precipitation*, *days with precipitation higher than 10 mm*, *summer soil moisture* and the *control of erosion rates* were not significant (p = 0.2), (p = 0.2), and (p = 0.7) respectively (Fig 6c-i). Higher values of *soil erosion* and *loss of soil organic matter* occurred in ecosystems providing higher *control of erosion rates* (p < 0.05) (Fig 6j-k). In contrast, the *density of semi-natural areas* and *livestock density* were low where the *control of erosion rates* was high or very high (p < 0.05) (Fig 6m and q). Moreover, there was no significant relationship between *crop diversity*, *fallow land*, *arable land* and *soil organic carbon* with the *control of erosion rates* (p = 1) (Fig 6l, n, o-r). The condition indicators *soil erodibility*, *bulk density*, *soil organic carbon* and the ecosystem service indicators *soil erosion risk* and *provision capacity* showed a positive relationship with the *control of erosion rates* (p < 0.05) (Fig 6s-v).

#### 3.2.2. Overlaps between environmental pressures, ecosystem condition, soil erosion risk and ecosystem service provision capacity

Fig 7 shows the spatial distribution of environmental pressures and condition in relation to the ecosystem service *provision capacity* in Lower Saxony. Fig 7a shows the superimposition of the maps of the normalized values of condition and provision capacity. Areas with high provision capacity and high condition levels (darker colours on the right top corner) were mainly located in the northwestern and central regions. High provision capacity was also found in municipalities with medium condition levels located mainly in the southern and eastern regions (dark blue). High provision capacity was not necessarily associated with a high level of ecosystem condition in municipalities such as the Harz in the southeast (light green). However, it is important to highlight that this area is mostly covered by mountainous forest and is part of a national park. On the contrary, some municipalities in the northwest showed low provision capacity, but medium condition levels (light blue). No municipalities in Lower Saxony were found to have low provision capacity and low condition levels.

**Fig 7.**
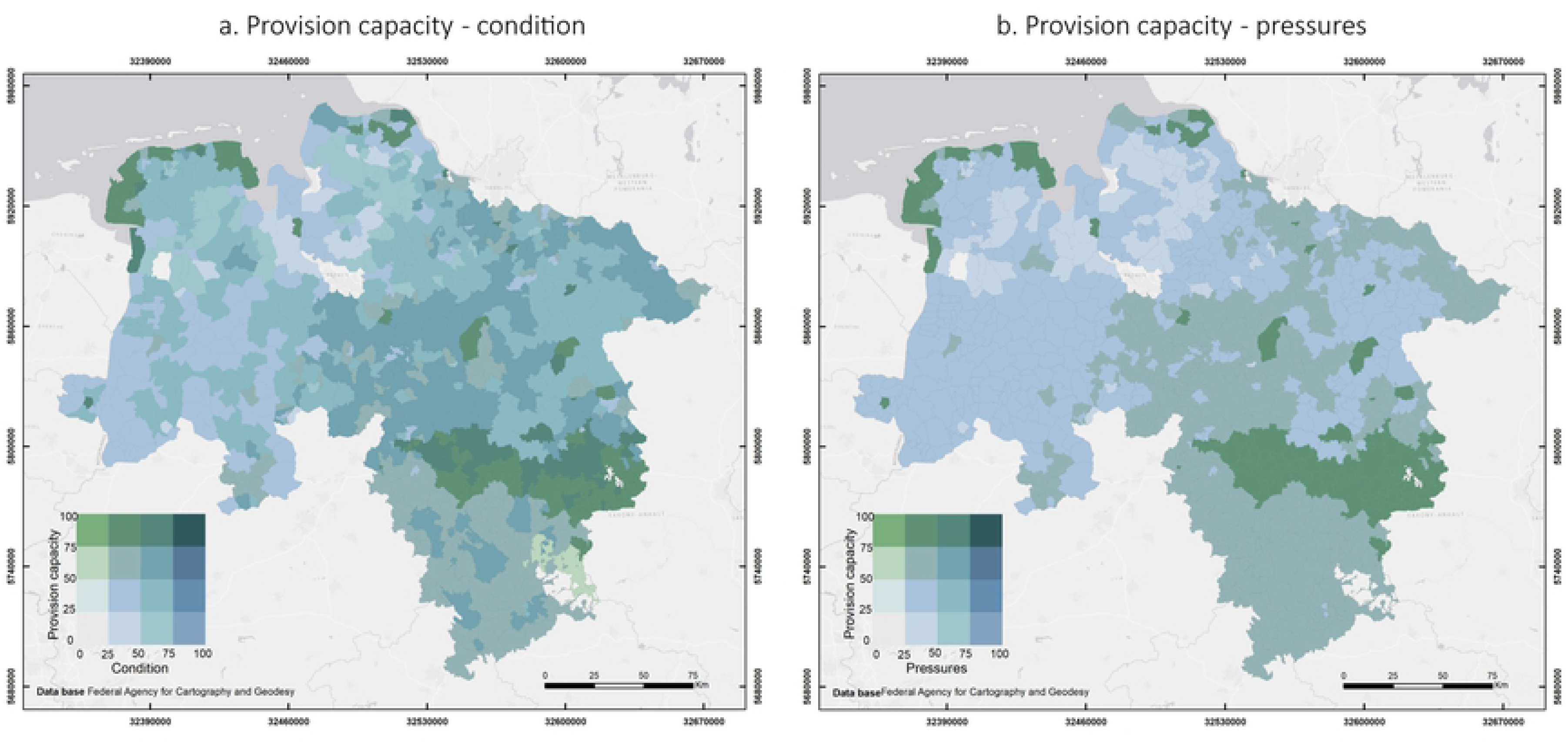
Spatial representation of the overlap between environmental pressures and condition, and ecosystem service provision capacity. (a) Overlap between provision capacity and condition. (b) Overlap between provision capacity and pressures. Darker areas represent high provision capacity and high level of condition in (a), or high provision capacity but high pressures in (b). Lighter areas represent lower provision capacity and low level of condition in (a) or lower provision capacity and low pressures in (b). (Large maps in Supplementary material S2).

The spatial distribution of provision capacity and pressures shows that most of the study area had medium levels of pressures and medium or high provision capacity (darker blue colours) (Fig 7b). The highest provision capacity occurred in the central and northwestern regions where the pressures had a medium level (dark green). Low provision capacities and medium pressure levels occurred in the northwest, especially in some municipalities of the districts of Cuxhaven, Rotemburg (Wümme) and Wesermarsch.

Fig 8 shows the spatial distribution of soil erosion risk, environmental pressures and condition in relation to the ecosystem service provision capacity in Lower Saxony. Fig 8a shows the superimposition of the maps of the normalized values of soil erosion risk, condition and provision capacity. Areas with high provision capacity, high condition levels and low erosion risk (top of the triangle) were mainly located in the northwestern region. High provision capacity was also found in municipalities with medium condition levels and low erosion risk located mainly in the western region and (to a lesser extent) in the northeastern (blue colour on the right side of the triangle) and the south (grey colour on the right side of the triangle). Medium provision capacity, condition and erosion risk were found in the district of Holzminden in the south (grey colour in the middle of the triangle). On the other hand, high provision capacity was evident in areas with medium-low condition levels and medium erosion risk (plum colours on the right side of the triangle). These mismatches were evident in the northwestern and western regions. Medium provision capacity, low condition levels and high erosion risk (tan colour at the bottom of the triangle) were evident in the southern region, especially in the municipality of Wenzen, also in the district of Holzmiden.

**Fig 8.**
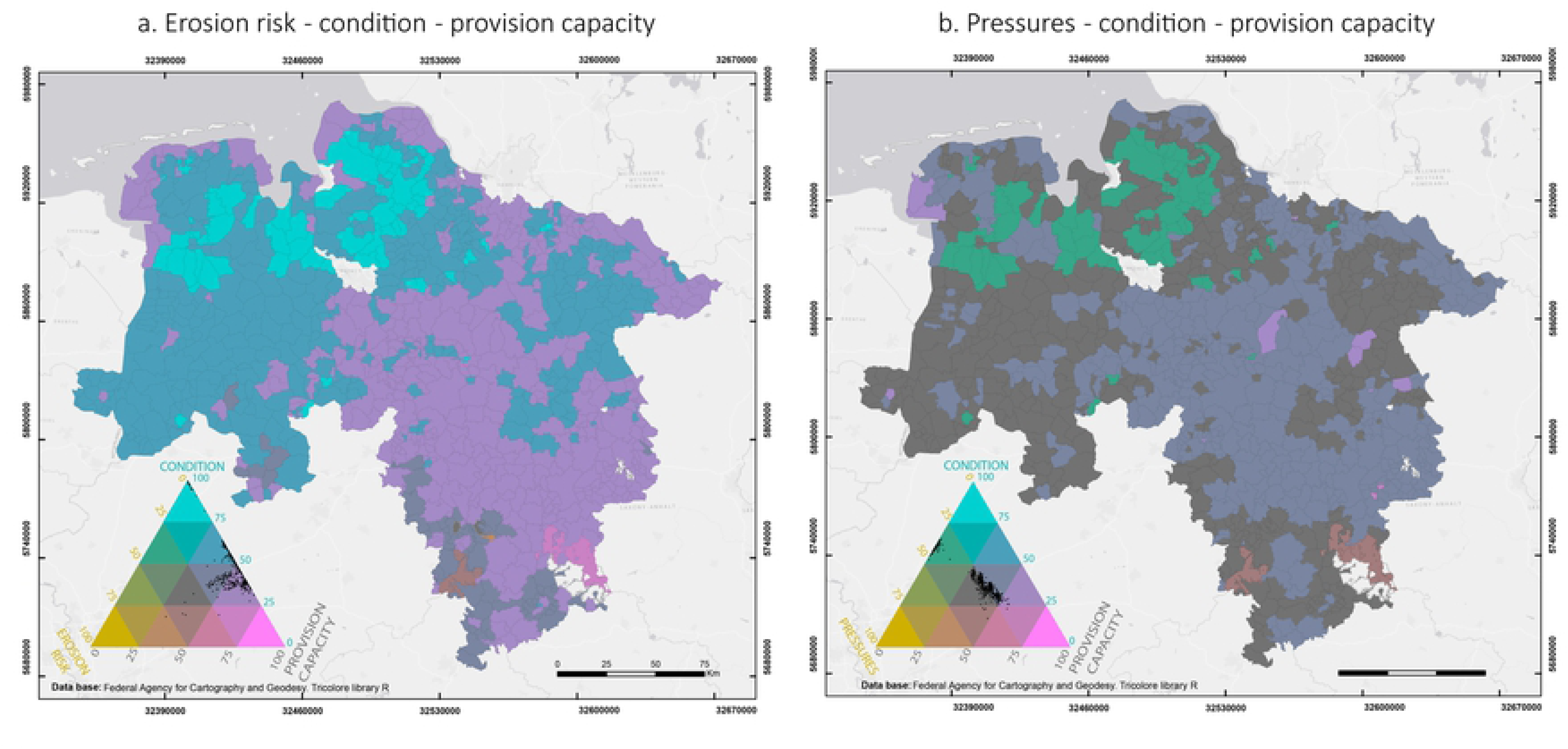
Spatial representation of the overlap between environmental pressures, condition, soil erosion risk and ecosystem service provision capacity. (a) Overlap between soil erosion risk, condition and provision capacity. (b) Overlap between pressures, condition and provision capacity. The colours and dots in the triangle show the distribution of the municipalities across the different indicators in percentage. Darker areas represent medium values of all the variables. Dots close the lower right corner indicate high values of provision capacity. Dots close to the lower left corner indicate high erosion risk in (a) and high pressures in (b). Dots close to the upper corner indicate high condition levels. (Large maps in Supplementary material S2).

The spatial distribution of provision capacity, pressures and condition shows that most of the study area had medium levels of pressures, medium condition levels and medium or high provision capacity (grey colours in the middle and on the right side of the triangle) (Fig 8b). The highest provision capacity was evident in the districts of Northeim and Goslar, but the condition levels in these areas were low and the pressures were medium. The lowest provision capacity occurred in the northwestern region where the pressures had medium levels and condition had high levels (green colour on the left side of the triangle).

## 4. Discussion

The analysis based on the operational MAES framework proposed by Maes et al [12] and the complied maps provide good results in regard to the relationships between ecosystem condition and soil erosion regulating ecosystem service in Northern Germany. In the following, we will discuss the limitations of the applied indicators and provide recommendations for improved applications.

### 4.1. Limitations of the indicators for ecosystem condition and control of erosion rates

#### 4.1.1. Environmental pressures indicators

*Change in ecosystem extent* was calculated at the municipality level, between the years 2016 and 2017 since there was no official statistical data available at this level for other time periods. This poses some limitations when assessing trends, because two years do not provide enough information to draw clear conclusions. Additionally, the indicator, as it is proposed here, only shows the degree of change, but not the drivers of change, so when it comes to comparison with an ecosystem service such as *control of erosion rates*, the mere percentage does not indicate whether the change is positive or negative for the provision of the service.

The analysis of the causes of change is important to understand the relationships between ecosystem condition and ecosystem services, but it was beyond the scope of this study, that aimed at testing the feasibility of the already proposed indicators (from the MAES working group) to assess ecosystem condition and to relate them to ecosystem services.

*Climate parameters* were calculated based on climate data for Germany for at least 30 years whenever they were available. However, there are some uncertainties with these data, and also other data used in the study, caused by the different methods of obtaining them as well as interpolations and missing or erroneous observations. In addition, for the climatic variables, the measurement network has changed over time and this affects the comparison of grid fields for the different years [45]. Another possible limitation of the indicator *climate parameters* is that the MAES framework does not provide guidelines about which specific parameters should be used to assess climate change. For this reason, we selected the most relevant indicators based on the possible influence on the ecosystem service *control of erosion rates*, which may seem arbitrary when looking at the general condition of the ecosystem or when making comparisons with other ecosystem services other than control of soil erosion, but a valid approach to generate data on soil erosion rates on the regional scale of this case study.

*Soil erosion* was calculated for arable land only, without taking grasslands and forests into account. Additionally, this calculation was made only for water erosion, without including wind erosion, which is a major problem in the northern part of Lower Saxony, therefore, the actual soil erosion might be higher than the results showed in this study.

Moreover, the calculation of soil erosion was made using the USLE equation that has some limitations, including the underestimation of the impact in thalwegs and gully erosion [43]. Furthermore, we used agricultural statistics and common assumptions about the effects of management practices and conservation measures for the estimation of soil erosion [48] which is less precise and less explicit than using detailed monitoring data [25].

*Loss of organic matter* values were obtained from the ESDAC, which was considered a useful and sufficient approach for our study area. However, as these values were originally calculated for the continental scale of Europe, some uncertainties exist regarding the accuracy of the results for the regional scale. Although the application of the CENTURY model at regional levels is technically feasible, there is still a lack of input data [50]. The collection and processing of these data would help to improve the accuracy of the results. However, it is very time-consuming and out of the scope of this study.

#### 4.1.2. Ecosystem condition indicators

*Crop diversity* was estimated based on the number of crop species in a diameter of 10 km (see Section 2.7.2). Although this calculation differs from the one suggested in the MAES framework (N° of crops/10 km × 10 km), it is also an approximation that reflects crop diversity. However, similar to the indicator *change in ecosystem extent*, this indicator does not provide sufficient information to determine whether this number is favourable or not when comparing it with the ecosystem service *control of erosion rates*. For this, an indicator such as the types of crops in a specific area would be more informative, since some cultures are more prone to soil erosion than others. However, it was not calculated in this study, since our aim was to test the feasibility of the proposed indicators.

*Density of semi-natural areas* was calculated in regard to the area of each municipality and not specifically in regards to the area of the agricultural land. This could overshadow the results since we did not determine the exact location of these elements, but instead estimated their proportion within the municipalities. Additionally, the lack of suitable metrics prevented us from calculating the shares of some semi-natural elements such as semi-natural grasslands, hedgerows and buffer strips that could have provided a different picture when it comes to identifying the role of semi-natural vegetation in the provision of ecosystem services such as *control of erosion rates*.

The indicators *share of fallow land, share of arable land and share of permanent crops in utilized agricultural areas* as well as *livestock density* were calculated based on official statistical data at the level of the municipalities. However, important factors such as the duration and the management of the fallow land can show different results regarding soil erosion in comparison to cultivated lands [72]. Nonetheless, the results could have been more precise and comparable with other indicators, if spatially explicit data were available for the study area. Furthermore, the bare number of livestock units per hectare does not provide sufficient information about the possible impact of livestock on the provision of ecosystem services, because the different types of livestock as well as their management (e.g. landless production systems vs grassland-based) are not assessed by this indicator.

*Soil Organic Carbon* was calculated based on spatially explicit data on humus content in the topsoil and then using common conversion factors to obtain the concentration of soil organic carbon. These spatially explicit results were upscaled to the level of the municipalities in order to allow for the comparison between indicators. However, as with other indicators, this leads to uncertainties in the results, since the soil characteristics, in this case soil organic carbon, are not homogeneously distributed within the area of the municipalities.

*Soil erodibility* or K factor of the USLE showed the same limitations as the calculated soil erosion described before. Furthermore, upscaling these values to the level of the municipalities can increase uncertainty, taking into consideration that the contents of silt, sand, clay and organic matter, as well as other parameters needed to calculate the K factor, usually vary within short distances and these generalizations can lead to less accurate results.

*Bulk density* values were obtained from the ESDAC, which was considered a useful and sufficient approach for our study, as well as for the indicator loss of organic matter mentioned before. However, bulk density shows limitations similar to the indicator loss of organic carbon as these values were also originally calculated for a continental scale. The processing of available regional data would improve the accuracy of the results, but this would be a very complex and time-consuming approach. It is also important to take into account the high annual variability of this indicator, which can be another source of uncertainty in this type of studies.

#### 4.1.3. Ecosystem service indicators

The ecosystem service indicator *potential soil loss* (erosion risk) was calculated assuming that the whole arable land is bare soil, taking into account only natural soil erosion by water. The results aggregated to the municipalities provide an approximate risk of erosion and could show a general picture of the ecosystem condition, when combined with other indicators.

As mentioned in Section 3.1.3, low values of *prevented soil loss* occurred also in areas with low *soil erosion risk*, but this does not necessarily mean that the service supply is low. This highlights that only calculating the *prevented soil loss* is insufficient to determine the actual ecosystem service supply [25].

The *provision capacity* indicator, which reflects the proportion of the potential soil loss that is mitigated by the ecosystem service *control of erosion rates*, allows to identify the service supply and to possibly assess different management practices. However, as with other indicators, provision capacity was upscaled to the municipality level and some aspects - e.g. the presence and distance to water courses, relief characteristics like thalwegs, presence of tramlines and wheel tracks, as well as management measures, which affect the provision capacity [24, 73] - could not be identified and hence the results are less accurate than they would be with spatially explicit data.

### 4.2. Relationships between agroecosystem condition and control of erosion rates

Since the adoption of the EU Biodiversity Strategy to 2020, the mapping and assessment of European ecosystems and their services has been increasing [74–76]. However, understanding the interdependencies between biodiversity, ecosystem functioning and ecosystem services is still a major challenge [5, 6]. Although, several studies provide evidence of the positive relationships between biodiversity, natural capital and ecosystem services [2, 77], there is no consensus on what these links are and how they concretely operate [78].

Although we were not able to establish the causalities among the indicators with the correlation analysis, we could observe that some indicators are strongly related. Almost all the environmental pressure indicators are strongly related to the ecosystem service *control of erosion rates*, except for the indicators *change in ecosystem extent*, *mean annual precipitation*, *days with precipitation equal or higher than 10 mm* and *summer soil moisture*. Regarding the ecosystem condition indicators, we identified that five out of nine condition indicators are strongly related to control of erosion rates. The four indicators that do not show a strong correlation are *crop diversity*, *share of fallow and arable land*, and *soil organic carbon*. As expected, there is a positive correlation between the ecosystem service indicators *potential soil loss* and *provision capacity* and the *control of erosion rates*. It must be noted that this analysis must be perceived with caution, considering the limitations of each of the indicators mentioned above.

### 4.3. Recommended land management measures to reduce soil erosion

Based on the overlaps presented in Section 3.2.2., we identified municipalities as priority areas in which the risk of soil erosion is medium or high, the provision capacity is low and the condition levels are low. Our data enable the identification of these *problematic* municipalities located in the district of Northeim in the southern region of Lower Saxony. For these areas, it is important to implement measures to reduce the impact of pressures, improve the ecosystem condition and soil conservation. In addition, analyses on local scale must be carried out and measures must take into account site-specific characteristics such as soil, crop varieties, soil degradation and farming practices [19]. When looking at the types of crops in Northeim, we identified cereals, grasslands, oilseeds, pastures for extensive grazing, and fodder plants (maize) as the main agricultural land uses. These crops may certainly have effects on soil degradation and erosion problems due to excessive tillage and crop residue removal [43]. Therefore, the implementation of conservation farming is essential to solve these problems because it can reduce soil erosion by ensuring the protection of the soil surface with residue retention and increased water infiltration.

The techniques applied in conservation farming include permanent soil cover with crop residues, which protects the ground surface and provides organic material, thereby improving soil quality. Other techniques are the growth of diverse crop species in the same field and crop rotation, especially crops such as legumes and grasses [34]. Conservation farming also involves minimum soil disturbance that has positive effects on biotic soil activity and leads to increased stability of soil aggregates [35]. Another practice that could be applied to guarantee soil conservation in vulnerable areas is agroforestry. This practice consists of the integration of trees with animals or crops or both, in order to preserve the fertility and structure of the soil through the fixation of nitrogen and the return of organic matter to the soil [34]. All these measures have an impact on the condition of soils and hence in the condition of agroecosystems, therefore, indicators that address these measures should be taken into account. The indicators proposed by the MAES working group, as they are implemented in this study, are not able to fully address agroecosystem condition relevant for soil protection and soil erosion prevention. Indicators that analyse the effect of crop species on soil erosion, the use of cover crops and other soil conservation measures should be included in the list of proposed indicators.

### 4.4. Potential for MAES/policy implementation

The normalization presented in section 2.9 and Figs 7 and 8 aimed to facilitate the comparison between indicators and to visualize the relations between ecosystem condition, environmental pressures, erosion risk and provision capacity. It is important to highlight that these representations are somewhat arbitrary and possibly other combinations and overlaps of indicators could provide different results. Furthermore, the results presented here come from a methodological study, aiming at testing an existing framework and respective indicators.

These results do not yet have the potential to be used to make policy decisions or implement the measures described above to improve the condition or reduce the pressures on agroecosystems. At least not without more detailed, and if possible, spatially explicit data, including more complete information on types and intensities of management practices. The maps presented in this study are intended to provide a general idea of the environmental pressures, ecosystem condition, soil erosion risk and the provision capacity of the ecosystem service *control of erosion rates* in Lower Saxony and to raise awareness of areas where special attention should be paid to avoid or mitigate ecosystem degradation.

Composite indicators can be used to provide insights on environmental condition, as well as sustainability, quality of life, and economy [79]. Such indicators have been useful in policy analysis and public communication because they seem to be easier for the general public to interpret [80]. However, we did not develop a composite indicator for three main reasons: First, it could add an extra ambiguity to the results. Second, because the suggested indicators would not be sufficient to build a trustworthy index, since threshold levels that would help to determine the overall condition have not been defined for all the indicators. Third, it was not the aim of the study to come up with a composite indicator, but to test the ones already proposed.

## 5. Conclusions

This is the first study testing the MAES framework and indicators for the assessment of the condition of agroecosystems in a regional scale case study. Furthermore, it is also an analysis of the relationships between ecosystem condition and the provision of a selected ecosystem service, specifically *control of erosion rates*. This assessment is useful to identify the suitability of these indicators, check the data availability for respective indicator quantification and to describe ecosystem condition in a regional scale.

Although we were not able to establish clear causalities among the indicators, our results identified positive, negative and no significant correlations between the different pressures and condition indicators, and *control of erosion rates* despite the limitations of the indicators and data availability. The idea behind the MAES framework is to show the general condition of an ecosystem in the context of ecosystem services supply. However, when looking at the relationships between ecosystem condition and ecosystem services, we identified that not all proposed indicators are suitable to explain to what extent agroecosystems are able to provide certain ecosystem services. Condition indicators on crop management and soil conservation measures, which are directly linked to the ecosystem service *control of erosion rates*, are missing in the list of indicators proposed in the MAES framework. Additionally, if indicators are to be applied in national or regional scale studies, it is important to consider that trend and high resolution data are not always available in sufficient quality and resolution, which may undermine the results and hence their comparability with other regions.

A further aspect to consider in future research is the assessment of other ecosystem services/ecosystem services bundles provided by agroecosystems. This would facilitate the identification of synergies and trade-offs, both between ecosystem services and between ecosystem condition parameters that may have a different degree of influence on ecosystem services. Moreover, a clearer definition of reference conditions, although complicated for agroecosystems, is important to provide more accurate information on the condition of the ecosystem, which should lead to better policy and management decisions.

## Acknowledgements

This study is partly based on the outcomes of the Lower Saxonian soil erosion monitoring programme funded by the Lower Saxonian State Authority for Mining, Energy and Geology (LBEG). We wish to thank the staff of the Joint Research Centre of the European Commission who gave conceptual support. We also wish to thank Tobias Kreklow and Jan Bierwirth for their collaboration in processing the data and Angie Faust for checking the language of the manuscript.

## Supplementary material

**S1. Indicators per municipality.**

**S2. Maps.**

**S3. Results of tests per indicator.**

